# Transdermal Delivery of Ultradeformable Cationic Liposomes Complexed with miR211-5p (UCL-211) Stabilizes BRAFV600E+ Melanocytic Nevi

**DOI:** 10.1101/2024.10.17.618694

**Authors:** Tanya Chhibber, Michael T. Scherzer, Anastasia Prokofyeva, Carly Becker, Rebecca Goldstein Zitnay, Eric Smith, Nitish Khurana, Mikhail Skliar, Dekker C. Deacon, Matthew W. VanBrocklin, Hamidreza Ghandehari, Robert L. Judson-Torres, Paris Jafari

## Abstract

Small non-coding RNAs (e.g. siRNA, miRNA) are involved in a variety of melanocyte-associated skin conditions and act as drivers for alterations in gene expression within melanocytes. These molecular changes can potentially affect the cellular stability of melanocytes and promote their oncogenic transformation. Thus, small RNAs can be considered as therapeutic targets for these conditions, however, their topical delivery to the melanocytes through the epidermal barrier is challenging. We synthesized and extensively evaluated ultradeformable cationic liposome (UCLs) carriers complexed with synthetic microRNAs (miR211-5p; UCL-211) for transdermal delivery to melanocytes. UCL-211 complexes were characterized for their physicochemical properties, encapsulation efficiency, and deformability, which revealed a significant advantage over conventional liposomal carriers. Increased expression of miR211-5p stabilizes melanocytic nevi and keeps them in growth-arrested state. We did a comprehensive assessment of cellular delivery, and biological activity of the miR211-5p carried by UCL-211 *in vitro* and their permeation through the epidermis of intact skin using *ex vivo* human skin tissue explants. We also demonstrated, *in vivo*, that topical delivery of miR211-5p by UCL-211 stabilized BRAFV600E+ nevi melanocytes in a benign nevi state.

## 1. Introduction

Melanocytes regulate pigmentation by producing melanin and are involved in a variety of skin conditions, including hyper- or hypo-pigmentation disorders such as melasma, chemical/drug-induced pigmentation, albinism, vitiligo, and melanoma [1–3]. Melanocytes are neural-crest-derived cells primarily located at the epidermal-dermal junction of the skin and are also distributed in hair follicles, inner ear, mucosa, iris (eye), and various organs throughout the body [2, 4]. The growth of melanocytes leads to the development of a variety of skin tumors, including benign lesions called melanocytic nevi (colloquially referred to as moles) and melanoma, one of the most dangerous forms of skin cancers [4–7]. Most of these diseases and conditions are driven by or associated with alterations in the expression of both coding, and/or non-coding regulatory genes [8–13]. The most common tumorigenic driver mutation in melanoma is the V600E mutation in the BRAF serine/threonine protein kinase, which drives over 80% of benign melanocytic nevi [4, 8, 9]. Melanocytes with BRAFV600E mutation proliferate to form nests of melanocytic nevi, which are growth-arrested.

MicroRNAs (miRNAs) are small, endogenous, non-coding RNAs that are powerful epigenetic regulators [14]. They form ribonucleoprotein RNA-induced silencing complex (RISC) in the cytoplasm that binds to specific protein-coding mRNAs, causing their degradation or inhibiting their translation [14, 15]. miRNAs play a major role in the regulation of important cellular processes such as proliferation, differentiation, or survival [16], and their dysregulation can lead to tumor initiation and progression by impacting various hallmarks of cancer [17, 18]. Due to their role in diseases, miRNAs have been explored as therapeutics (miRNA mimics or inhibitors), and as potential biomarkers for cancer diagnosis and prognosis [12, 16, 17, 19–21]. We have previously reported that miR211-5p is the most highly and differentially expressed transcript enriched in nevi, as compared to healthy skin melanocytes and other melanocytic tumors [6, 12, 16, 21], and is both necessary and sufficient to stabilize BRAFV600E+ melanocytes into a nevus-like state *in vitro* [6, 16]. This suggests that the overexpression of miR211-5p in BRAFV600E+ melanocytes *in vivo* would also stabilize the nevi.

Systemic delivery of miRNAs is associated with various challenges, including enzymatic degradation during circulation, decreased bioavailability at the target site, systemic toxicity, and rapid clearance [22]. Topical applications would overcome the systemic challenges and provide localized delivery to the target site. However, transdermal delivery of miRNAs to melanocytes also poses a challenge due to the barrier properties of the skin, low cellular uptake due to the negative charge of miRNAs, and degradation by nucleases [23]. Various groups have used microneedles [24] or intradermal injection-based [25] methods to deliver miRNAs or drugs to melanocytes. However, these techniques are somewhat invasive.

Multiple reports have highlighted the potential of Ultradeformable Cationic Liposomes (UCLs) to deliver small RNAs into deep epidermal layers including melanocytes [26–29]. However, to our knowledge, reports on direct biological responses triggered by the UCL-delivered small RNAs in skin melanocytes in *in vivo* models have not been reported. UCLs or elastic liposomes, differ from traditional liposomal systems as they consist of edge-activating agents and lipids, providing these vesicles with high elasticity and flexibility allowing them to cross the intact skin more efficiently and reach the deeper skin layers [26, 30–32]. The edge activating agents are single-chain surfactant molecules like sodium deoxycholate, sodium cholate, Span 80, and Tween 80, that destabilize the lipid bilayer of the UCLs, making the membrane more flexible compared to the conventional liposomes[26, 31, 33, 34]. The cationic nature of the UCLs makes it easy to electrostatically complex with negatively-charged RNAs and facilitate their delivery [27, 30]. Previous studies have demonstrated that the transport of UCLs through the skin is driven by differences in the hydration gradient in the skin layers, creating a strong force driving the UCLs to the deeper layers of the skin [30, 35, 36].

Here, we report on the synthesis and extensive characterization of the UCLs complexed with miR211-5p (UCL-211) and demonstrate successful delivery of biologically active miR211-5p *in vitro,* and UCL-211 permeation into skin layers *ex vivo.* Also, we show that topically applied UCL-211 successfully delivered biologically functional miR211-5p to the fields of melanocytes harboring BRAFV600E mutation in the skin overcoming the skin barrier, in a mouse model of nevus formation. The UCL-211-mediated overexpression of miR211-5p in BRAFV600E+ melanocytes led to the formation of melanocytic nevi in these fields, confirming the targeted delivery of the miR211-5p, and the maintenance of its biological activity.

## 2. Materials and methods

### 2.1 Materials

1,2-Dioleoyl-3-trimethylammonium propane methyl sulfate (DOTAP) and F-DOTAP (1-oleoyl-2-[6-[(7-nitro-2-1,3-benzoxadiazol-4-yl)amino]hexanoyl]-3-trimethylammonium propane (chloride salt)) were purchased from Avanti Polar Lipids, Inc. (Alabaster, Alabama, US). Sodium cholate hydrate (NaChol), Cholesterol and HEPES buffer 1M were purchased from Sigma-Aldrich (St. Louis, MO, US).

### 2.2 miRNA

The following miRIDIAN microRNA Mimics were purchased from Horizon Discovery Ltd (US). hsa-miR-211-5p (Human; Sequence-UUCCCUUUGUCAUCCUUCGCCU) mmu-miR-211-5p (Mouse; Sequence-UUCCCUUUGUCAUCCUUUGCCU) microRNA Mimic Negative Control #1 (mature sequence: UCACAACCUCCUAGAAAGAGUAGA A fluorescently (Cy5) tagged Custom miRNA Mimic for miR211-5p (miR211-5p-Cy5) was designed with the following details and purchased from Horizon Discovery Ltd. (Active: 5’-PUUCCCUUUGUCAUCCUUUGCCU3’ Passenger: 5’-**Cy5**GCAAAGGUGACAAAGGGAAUU3’).

### 2.3 Synthesis of UCLs

UCLs were synthesized by thin film hydration technique as previously described [27]. In brief, DOTAP at the concentration of 10 mg/ml was dissolved in chloroform, and sodium cholate (NaChol) at the concentration of 10 mg/ml was dissolved in ethanol, and the two solutions were mixed. Initially, UCLs at four different weight: weight ratios (w/w) of DOTAP: NaChol (4:1, 6:1, 8:1, and 10:1) were synthesized. Next, the organic solvents were evaporated using a rotary evaporator (BUCHI R210 rotavapor, BUCHI Corporation, New Castle, USA). The resulting thin film was kept overnight in a vacuum chamber to remove any residual organic solvents. The lipids were hydrated with 20 mM HEPES buffer at pH 7.4 and vortexed for liposome formation. The liposomes were then collected and stored overnight at 4°C. The next day, the liposomes were sonicated for 50 mins and then extruded 11 times through a 200 nm pore-sized membrane, followed by a 100 nm pore-size membrane using Liposome Hand Extruder (Genizer LLC, US). The extruded liposomes were then dialyzed using 50kDa membranes for 24 h to remove any free lipids from UCLs.

Fluorescently labeled liposomes were prepared by the same method described above with the addition of green fluorescently labeled DOTAP (F-DOTAP-consisting of NBD (nitrobenzofurazan) tag conjugated to DOTAP) at a 100:1 molar ratio (DOTAP: F-DOTAP) to the reaction.

### 2.4 Complexation of miR211 with the UCLs

To load the miR211-5p mimic onto empty UCLs, the mimic was dissolved in nuclease-free water and passively complexed with the liposomes at different nitrogen: phosphate (N/P) ratios to obtain UCL-211. The N/P ratio was calculated using the equation below [37]. An N/P ratio of 4 (40 µM) was selected for this study.

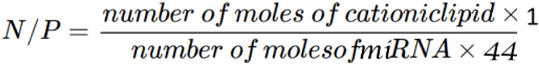

### 2.5 Measurement of Size, Size Distribution, and Zeta Potential

The average particle size, polydispersity index (PDI), and zeta potential of the synthesized UCL and UCL-211 were determined before and after complexation with miR211-5p, using a Zetasizer instrument (Malvern Panalytical Ltd, Westborough, Massachusetts, US). UCL and UCL-211 were diluted in Milli-Q water for characterization by dynamic light scattering (DLS) technique, and in 10 mM NaCl solution to determine the zeta potential. The average particle size, PDI and zeta potential were measured for samples from three independent batches, with 14-16 measurements per sample.

We used Nanoparticle Tracking Analysis (NTA) to characterize the size distribution and concentration of background particles in buffer (HEPES Blank), UCL and UCL-211 samples [38]. First, liposome samples were diluted with Milli-Q water to bring the particle concentration into the recommended range [39]. Samples were then flown through the test chamber by an automated syringe pump (Harvard Bioscience, MA) and illuminated by a red laser (642 nm wavelength; Nanosight model LM14, Malvern, Salisbury, UK). The light scattered by individual particles was video recorded for 60 seconds by a high sensitivity sCMOS camera (Hamamatsu Photonics, Japan) using a 32.5 ms shutter speed and 25 fps (frames per second) acquisition rate. The measurements were repeated five times for each sample, keeping the camera level at 16. Recorded videos were analyzed by NTA software (Nanosight version 3.4) to extract particle tracks and calculate mean-squared particle displacements from the bulk flow and the corresponding diffusion coefficient. The hydrodynamic diameter of each particle was then obtained from the Stokes-Einstein relationship using the measured temperature and assuming the samples have the same viscosity as DI water [39, 40]. The particle detection threshold during the analysis was set to 10, and the minimum length of particle tracks to be included in the analysis was selected as Auto. The data for all particles imaged during repeated measurements were separately averaged for each sample to obtain the particle size distribution and concentration. Standard deviations from the mean were used to quantify the uncertainty of the obtained results.

### 2.6 Gel Retardation Assay

The optimal complexation of the empty UCLs with miR211-5p mimic was assessed by performing a gel retardation assay that evaluates the charge neutralization due to the complexation of negatively charged miR211-5p mimics with positively charged UCLs. Free miR211-5p mimic and UCL-211 at different ratios of miR211-5p: UCL (1:1, 1:2, 1:4, 1:8, 1:12, and 1:16) were loaded and gel electrophoresis was performed on 4% agarose gel with 0.5% ethidium bromide, and the gel was visualized under UV light. We used the 1:1 batch as a negative control [27].

### 2.7 Colloidal Stability

The colloidal stability of UCLs was assessed by measuring the particle size over a period of one-month post-synthesis at the standard storage temperature of 4° C. The stability of UCL-211 was determined by particle size and by visual inspection for the absence of aggregates immediately after complexation.

### 2.8 Morphology

The morphology of UCL and UCL-211 liposomes were visualized using Transmission Electron Microscopy (TEM). The UCL samples were loaded onto ultrathin holey carbon films supported by 400 mesh copper grids and air-dried in a dust-free environment. The grids were treated under plasmonic discharge and then observed using TEM (JEOL 2800 TEM/STEM; JEOL USA, Inc., Peabody, MA, US, at the Nanofab facility at the University of Utah).

### 2.9 Entrapment efficiency

The amount of miR211-5p complexed with the UCLs was calculated using Quant-iT™ microRNA Assay Kit (Invitrogen™, US). A method previously described with some modifications was used [41, 42]. The samples were prepared by diluting UCL-211 in 1X TE buffer with 0.5% Triton-X 100, sonicated for 10 min at 37 °C, and then incubated for an hour. UCL-211 was diluted in 1X TE buffer without Triton-X 100 as a control. The samples were prepared according to the manufacturer’s protocol and analyzed by measuring fluorescence at excitation/emission maxima of 500/525 nm. The encapsulation efficiency was calculated using the equation-

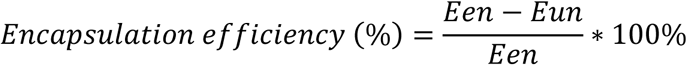

Where Een was the total amount of miR211-5p, and Eun was the amount of free miR211-5p (in 1X TE buffer) [42, 43].

### 2.10 Lipid quantification

Stewart assay was used to quantify the lipid component of UCLs based on the previously described method with slight modifications [44, 45]. In brief, ferrothiocyanate reagent was prepared using ferric chloride hexahydrate and ammonium thiocyanate in double distilled water. 200µL of the sample was mixed with 200uL of chloroform and vortexed for 20 mins. 400µL of prepared ferrothiocyanate reagent was added, and the tubes were again vortexed for 20 mins. The tubes were centrifuged at 10,000 RPM for 20 mins. The lower chloroform layer was quickly removed (to minimize evaporation of chloroform) and the absorbance was read at 485nm. Initially, a standard curve for DOTAP was prepared (with a dilution factor of 2-10 mg/mL-0.625 mg/mL) and the equation obtained from the standard curve was utilized for determining the lipid content in the samples.

### 2.11 Deformability analysis

To assess deformability, the elasticity of UCLs and conventional non-deformable liposomes (NDLs) was compared using a previously described method with minor modifications [46]. NDLs were synthesized by the same method as described for UCLs, with modifications in the lipid composition. For NDLs, we replaced NaChol (edge activating agent responsible for providing the deformability characteristic), with cholesterol. One mL of UCLs and NDLs were extruded through a membrane with a pore size of 50 nm (25 mm diameter) at 100 psi pressure (0.68 mPa). The amount of extruded DOTAP was calculated using the Stewart assay for lipid quantification (Section 2.10). Three measurements were taken on two batches of UCL and NDL formulations. Equations previously described in [46] were used to measure the elasticity of the UCLs.

### 2.12 Cell Culture

Human Epidermal Keratinocytes (HEK), Human Dermal Fibroblasts (HDF), and Normal Human Melanocytes (NHM), medium-pigmented and dark-pigmented melanocytes were used. Cells were isolated from fresh skin specimens as previously described [47]. All clinical specimens were collected with informed consent under a protocol approved by the Institutional Review Board (IRB_00010924), and the study was conducted in accordance with the ethical standards of the 1964 Helsinki Declaration and its later amendments. The cells were kept in controlled humid conditions in incubators at 37°C and 5% CO_2_. NHMs, and medium and dark pigmented melanocytes were grown in Gibco™ Medium 254 (Gibco™, US) supplemented with Gibco™ Human Melanocyte Growth Supplement-2 (HMGS-2; Gibco™, US-referred to as HMGS2 media) or Gibco™ Human Melanocyte Growth Supplement (HMGS; Gibco™ US-referred to as HMGS media). HEK were grown in the Keratinocyte Growth Medium 2 kit (PromoCell, Germany). HDFs were grown in Gibco™ DMEM (Gibco™, US) supplemented with 10% FBS (fetal bovine serum) and 2 mM Gibco™ L-Glutamine (Gibco™, US). During treatment with UCLs, the media was also supplemented with Gibco™ Antibiotic-Antimycotic (Gibco™, US).

### 2.13 Cytotoxicity studies

The cytotoxicity of UCL formulations was assessed on different primary human skin cell types, including NHM, HEK, and HDF, using Cell Counting Kit-8 (CCK-8) assay (Dojindo Molecular Technologies, Inc., US) following manufacturer protocol. The cells were grown in a 96-well plate overnight and treated with the formulations at a dilution of 1:8 in culture media for four hours. Then the cells were washed twice, and fresh media was added to each well. After 24 h, the CCK-8 assay was performed according to the manufacturer’s instructions, and the absorbance was measured at 450 nm using a UV spectrophotometer (Thermo Scientific Varioskan LUX Multimode Microplate Reader, US).

### 2.14 Uptake of UCLs by primary cells

Uptake of fluorescently labeled UCLs by primary NHM with different pigmentation levels (light, medium, and dark), HEK, and HDF was evaluated by confocal microscopy and flow cytometry. For imaging, cells were grown on crystal clear glass coverslips coated with poly-L-lysine (GG-12-PLL, Neuvitro Corporation, US), and treated with HEPES buffer 20mM (control), and fluorescently labeled UCL and UCL-211 (diluted in HEPES buffer 20mM) for 6h. The cells then were washed and fixed in 4% PFA (paraformaldehyde) for 15 minutes. Nuclei were counterstained, and coverslips were mounted on slides using ProLong™ Glass Antifade Mountant with NucBlue™ Stain (Invitrogen™) and imaged using a Nikon Ti-E inverted microscope (Nikon, US).

For flow cytometry, cells were treated with fluorescently labeled liposomes for 6 hours, washed, and collected. The cells were then analyzed using Beckman Coulter Cytoflex S (Beckman Coulter Life Sciences, US). Similar uptake studies were also conducted for NHM with different pigmentation levels (light, medium, and dark) in different media conditions (HMGS and HMGS2) to determine the effect of different levels of melanin pigment and melanocyte phenotypes on the uptake of UCL-211.

### 2.15 Quantification of miR211-5p uptake by primary cells

NHM cells were seeded at a density of 2×10^5^ cells per well in a 12-well plate and grown in HMGS2 media. The cells were treated with either HEPES buffer 20mM (control), or UCL-211 and UCL-NC (UCLs complexed with non-targeting control miRNA) for 6 h. Cells were then washed after 6 h, and fresh HMGS2 media was added to treated groups. A group of cells treated with HEPES buffer 20mM for 6h, were transitioned to HMGS media after the washing step. The cells were incubated for 72 h in HMGS2 or HMGS media and then detached and processed to perform absolute qPCR analysis. After 72 h, the cells were detached with Cell Dissociation Buffer Enzyme-free Hanks’-balanced salt solution (Gibco™, US) for 15-30 minutes, pelleted and resuspended in phosphate-buffered saline (PBS) supplemented with 0.5% bovine serum albumin and 2 mM EDTA (rinsing solution), re-pelleted, and resuspended in 300 µL of rinsing solution. After adding 100 µL of FcR blocking Reagent (Miltenyi Biotec, US), and 100 µL of CD117 MicroBeads suspension (Miltenyi Biotec, US), the cells were mixed and incubated for 15-30 min on ice. An LS column (Miltenyi Biotec, US) was fitted onto a “The Big Easy” EasySep Magnet (Stemcell Technologies, US) and activated by gravity filtering 3 ml of rinsing solution. The CD117 labelled cells were added to the column and rinsed 3x with 3 ml of rinsing solution by gravity filtration. The column was then removed from the magnet, 5 ml of rinsing solution was added to the column, and the cells were released using a plunger. Cell pellets and wash buffers underwent RNA extraction using the miRNeasy Micro Kit (Qiagen cat. No. 217084) according to the manufacturer’s protocol or the miRNeasy Serum/Plasma kit protocol (Qiagen cat. No. 217184), respectively. RNA was quantified using Qubit dsDNA Quantification, High Sensitivity Assay (Thermo Fisher cat. No. Q32851). miRNA reverse transcription and amplification were performed using TaqMan™ Advanced miRNA cDNA Synthesis Kit (Thermo Fisher cat. No. A28007) with 10 ng of RNA input per sample. Amplified miRNA cDNA samples were diluted 1:500 in 0.1X TE. miRNA specific Taqman Advanced miRNA assays (Thermo Fisher cat. No. A25576) were used to amplify miRNA of interest on the Applied Biosystems QuantStudio AbsoluteQ Digital PCR System using the AbsoluteQ DNA digital PCR Master Mix (Thermo Fisher cat. No. A52490) protocol. For each sample the analysis was performed using QuantStudio AbsoluteQ Software (v6.3) to generate concentration data (copies/µL) for the amplified hsa-miR-211-5p (Assay ID: 478507_miR, Thermo Fisher Scientific Inc., US) and the control hsa-miR-16-5p (Assay ID: 477860_miR, Thermo Fisher Scientific Inc., US). This experiment was repeated in three biological replicates.

### 2.16 Biological activity using a miRNA dual reporter assay

For this study, pLVX-anti-MIR211-5p (Addgene #153318; http://n2t.net/addgene:153318) previously developed by our group [6] was used to assess the activity of miR211-5p delivered by UCL-211.

#### Generation of lentiviral particles

Lentiviral transduction was done using HEK293T (human embryonic kidney cell line) cells on a 10cm tissue culture dish. The cells were plated at 1.8×10^6^ and transfected using the method previously described in [6] with jetPRIME transfection reagent (Polyplus, 712-60). We then collected and filtered the viral supernatant 48 h post-transfection, using 0.45 µm syringe filters. NHM seeded at a cell density of 4.0×10^5^ cells/well in a 10-cm tissue culture dish were incubated with the filtered viral supernatant with 10 µg/ml polybrene (Sigma-Aldrich, TR-1003). After 6h, the viral supernatant was removed, the cells were washed twice, and fresh NHM growth media was added. Transduced NHM cells were sorted for GFP and mCherry expression within the linear range of the reporter using a BD FACSAria II (BD Biosciences, CA).

#### MicroRNA reporter assay

We performed the reporter assay on the sorted population of cells expressing medium levels of both GFP and mCherry. Sorted cells were seeded in a 48-well plate at 1.5 ×10^4^ cells per well in HMGS2 media. The cells were treated for 6 h with either HEPES 20mM (control), UCL-NC, or UCL-211 formulations supplemented with HMGS2 media. Another control treated with HEPES 20mM was supplemented with HMGS media (positive control). After 6 h of incubation, the media was removed, and the cells were washed twice. Fresh HMGS2 or HMGS media was added to all cells. The plates were then placed on the Phasefocus Livecyte® microscope, and the cells were imaged every two hours for 72 h. Four different randomly chosen regions of interest in each well were live imaged using Livecyte Phasefocus Acquire software (v3.10.2) (Phasefocus, Sheffield, UK) and repeated in triplicate. The images were segmented and analyzed using ImageJ (FIJI) ratiometric analysis to determine GFP and mCherry levels. We used the phase images to form thresholds and then used the raw integrated intensity for the GFP and mCherry positive cells.

### 2.17 Evaluating morphological changes using Quantitative Phase Imaging (QPI) using Phasefocus Livecyte® microscope

Morphological changes of NHM cells following UCL treatment were assessed through Quantitative Phase Imaging (QPI) as following: NHM cells were seeded in 24 well plates at 1.5×10^4^ cells per well and grown in HMGS2 media condition. The cells were then treated with either HEPES buffer 20mM (negative control), UCL-NC, or UCL-211. After 6 h treatment, the cells were washed twice and fresh HMGS2 media was added to UCL-NC, UCL-211 and one group with HEPES buffer 20mM. The media of another HEPES buffer control group was changed to HMGS. The plates were then loaded on the Phasefocus Livecyte® microscope (PhaseFocus Limited, UK). Randomly chosen four different regions of interest in each well were live imaged every 2 hours for 72 h and analyzed using Livecyte Phasefocus Acquire software (v3.10.2) (Phasefocus, Sheffield, UK).

### 2.18 *Ex vivo* skin permeation

#### Franz diffusion assembly set up and skin preparation

Fresh full thickness deidentified human skin tissue was obtained as surgical discards from the Huntsman Cancer Institute (HCI) Biorepository and Molecular Pathology shared resource. All clinical specimens were collected with informed consent under a protocol approved by the Institutional Review Board (IRB_00010924), and the study was conducted in accordance with the ethical standards of the 1964 Helsinki Declaration and its later amendments. The skin was trimmed to remove most of the subcutaneous tissue, and appropriate sections were cut to fit the Franz diffusion cell dimensions. The tissue was then carefully mounted on the receptor chamber of the Franz diffusion cell with the stratum corneum side facing the donor compartment. The donor and receptor compartments were fixed with a clip to avoid any disruption during the process. The receptor compartment was filled with 20mM HEPES buffer and then placed in a Franz diffusion chamber at 37.0°C, with continuous stirring with a magnetic bar placed in the receptor compartment. 100 µL of each formulation was applied to the skin and spread evenly, which included control (20mM HEPES buffer), and fluorescently labeled UCL-211. The experiment was performed for 24 h. After 24 h, the skin was collected, and tissue fixation was performed.

#### Tissue fixation and preparation

The collected skin tissue from the Franz diffusion cell was fixed in 10% formalin for three days and then washed with 70% ethanol and PBS. After three days, the samples were washed with PBS. The tissue was submitted to ARUP Research Histology Core Laboratory (ARUP Laboratory, Salt Lake City) for paraffin embedding and processed for formalin-fixed paraffin-embedded (FFPE) sectioning. Slides with 5 µm sections of skin samples were obtained, and immunofluorescence staining was further performed.

#### Immunofluorescence staining for the skin tissue

The tissue slides were heated at 65° C, followed by deparaffinization steps and antigen retrieval using 10mM sodium citrate buffer, pH 6.0. The tissues were treated with the blocking buffer 1% bovine serum albumin (BSA), 5% normal donkey serum and 0.3% Tween-20 in 1X-DPBS for an hour followed by overnight incubation with primary antibody for melanocyte staining, Melan-A (1:100, Abcam ab731). The slides were washed the next day and incubated with a secondary antibody (Invitrogen Donkey anti-Goat IgG (H+L) Cross-Adsorbed Secondary Antibody, Alexa Fluor™ 594; A-11058) in the dark. The sections were counter-stained for nuclei and coverslips were mounted on slides using ProLong™ Glass Antifade Mountant with NucBlue™ Stain (Invitrogen™). Skin samples were imaged using a confocal fluorescent microscope (Leica SP8 405-488-561-633 Laser Confocal microscope-Leica Microsystems Inc., Deerfield, IL US).

### 2.19 *In vivo* transgenic mice model of BRAF^CA^/Tyr:CreER

All animal experiments were conducted in accordance with institutional guidelines and were approved by the Institutional Animal Care and Use Committee (IACUC, protocol 20-1200). Animal care and use procedures complied with the guidelines set forth by the National Institutes of Health and the Animal Welfare Act.

#### 2.19.a. Initiating Nevi with 4-hydroxytamoxifen (4-HT) in the Tyr::CreERT2 BRAF^CA^

Mice Cohorts of ∼6-8-week-old Tyr::CreERT2;BRAF^CA^ mice were bred [48, 49]. Prior to P17 (postnatal day 17), after the birth of pups, tail biopsies were performed which were further used to genotype all the pups using PCR to confirm they were BRAF^CA^: Tyr::CreERT2 positive. Genotyping was also done to segregate male and female pups. On P21, the mice were weaned off.

#### 2.19.b. 4-HT application to initiate BRAF^CA^ conversion to BRAF^V600E^

Six weeks after the birth of the pups, the back skin around the base of the tail was shaved ∼24 hours prior to applying 4-HT to initiate BRAF^CA^ conversion to BRAF^V600E^. Mice were anaesthetized under isoflurane during the 4HT application and until it dried to avoid spreading of 4-HT by self-grooming. 20µl of 25.8mM solution (10mg/1mL) of 98% 4-hydroxytamoxifen (Sigma # H7904-98% Z isomer) dissolved in 100% ethanol was applied on the shaved region. 4-HT was again applied the next day using similar procedure and the pups were then left to rest for three days.

#### 2.19.c. UCL Treatment

After three days, different formulations were applied to the region previously treated with 4-HT. Controls included mice with no 4-HT application and mice with 4-HT application. Treatment groups included Vemurafinib treatment (positive control), UCL-NC, UCL-211, NDL-211, and free miR211-5p (dissolved in HEPES buffer). Treatments were applied on alternate days for 21 days. On Day 21, the mice were euthanized by CO2 inhalation method (5-minute administration). The treatment area was waxed to remove any hair from the skin and the tissue was dissected. The skin was washed under running water, placed on a slide and imaged at 1X magnification using Olympus MVX10 microscope (Evident Scientific, Inc., US). The tissues were then fixed in 10% Neutral-Buffered Formalin. The images for the mice were then analyzed using a semi-quantitative scoring system. For each mouse, the four skin regions with the highest number of nevi were selected, and the number of nevi were counted. A nevus density scoring system was then applied. No-nevi regions were categorized as “0,” 1-50 distinct nevi were graded 1, 51-100 distinct nevi were graded 2, >100 nevi with generally distinct boarders were graded 3 and matts of nevi so dense that boarders could not be reliably assessed were graded 4. The average score of the four most nevus dense regions was calculated.

### 2.20 Statistical analyses

Statistical analyses were performed by calculating P values using one-way Anova using Dunnett’s multiple comparisons test or using paired or unpaired two-tailed t-tests via Prism 10 (GraphPad) as indicated in figure legends. Results were considered different with statistical significance if the p-value was <0.05. The data are represented as mean ± standard deviation (SD).

## 3. Results and Discussion

### 3.1 Synthesis and characterization of UCLs

Initially, UCLs were synthesized by thin film hydration technique with varying w/w ratios of DOTAP and NaChol ranging from 4:1 to 10:1 to determine the optimal formulation based on the following criteria: minimal cytotoxicity and maximal uptake by skin cells, and the highest permeability through the epidermis. Different UCL formulations were characterized by their particle size, polydispersity index (PDI), and zeta potential. The particle size was measured by Dynamic Light Scattering (DLS) for these UCLs and ranged between 85-95 nm (**Supplementary Figure S1.a**), with 1:4 ratio having the lowest particle size followed by 1:6 and 1:8 batches. Since the 1:4 batch had the highest NaChol along with DOTAP, these UCLs were expected to be more deformable which might have been a factor contributing to their smaller size. The PDI for all different ratios was less than 0.15 (**Supplementary Figure S1.a**). For stable and less polydisperse liposomes a PDI below 0.2 is considered ideal [50]. The zeta potential characterization was performed to determine the stability of these UCLs in buffer which was over +30mV for all the batches (**Supplementary Figure S1.b**). For a colloidal suspension to be considered stable, the zeta potential of the particles should generally be greater than +30mV or less than −30mV [51]. As DOTAP is a positively charged lipid, we expected to have positively charged UCLs. The zeta potential analysis confirmed that all four formulations were stable. These physical characteristics of UCLs play a crucial role in their ability to penetrate the epidermal skin layers and reach melanocytes located at the epidermal-dermal junction.

We then assessed the effect of these formulations on the viability of different skin cell types. We performed a CCK8 cytotoxicity assay on normal human NHM, HDF, and HEK cells, treated with these formulations for 4 hours in *in vitro* cultures. CCK8 assay is a highly sensitive colorimetric method for quantifying viable cells. None of the UCL formulations with varying DOTAP to NaChol ratio showed significant cytotoxicity in cells (**Supplementary Figures S1.c, S1.d, and S1.e**). We also assessed the cellular uptake of fluorescently labeled UCL 1:8 batch by NHM cells by imaging them after 4h of exposure. The cytoplasmic localization of green signal (corresponding to the NBD tag on the UCLs) detected by fluorescence microscopy of NHM cells incubated with UCLs, revealed the internalization of the UCL formulation. No fluorescent signal was observed in control cells treated with 20mM HEPES buffer (**Supplementary Figure S1.f**).

Next, we evaluated the skin permeability of fluorescently labeled UCL formulations using Franz diffusion chambers and deidentified human skin tissue biopsies. The skin was treated with fluorescently labeled UCLs for 24 h, followed by sectioning and immunostaining for melanocyte marker, Melan-A. Fluorescence microscopy of skin sections revealed that the UCLs with NaChol: DOTAP ratio of 1:8 could penetrate the skin layers and localize to epidermis (**Supplementary Figure S1.g; green signal**), while we did not observe any colocalization in the 1:4 and 1:6 batches. This was expected based on similar studies conducted previously [27]. As these formulations consisted of higher DOTAP concentration, it is possible the vesicles had a higher degree of deformability and crossed the skin layers over 24 h of incubation. We identified our optimal formulation based on the preliminary skin permeability studies to be UCLs with 8:1 DOTAP: NaChol ratio that demonstrated the best skin permeation characteristics. Therefore, we selected this formulation for loading miR211-5p mimic and further studies.

### 3.2 Complexation of selected UCL formulation with miR211-5p

The UCL formulation with a DOTAP: NaChol ratio of 8:1 was complexed with miR211-5p at different nitrogen/phosphorus (N/P) ratios. The phosphorous on the miRNA electrostatically complexes with nitrogen on the lipids to form UCL-211. Various N/P ratios from 1.6:1, 3:1, 4:1, and 8:1 were evaluated. The formulations with N/P ratio of 1.6:1 immediately crashed upon addition of miR211-5p. The 3:1 ratio was comparatively more stable but underwent precipitation over time. The N/P ratio of 4:1 and 8:1 did not show any visual aggregate formation upon the addition of miR211-5p to the UCLs. Since we wanted to load the maximum concentration of miR211-5p, we selected the N/P ratio of 4:1 (at 40 µM) for future studies. The complexed UCL-211 formulation with a UCL: miR211-5p ratio of 4:1 was then characterized for particle size, PDI, and zeta potential. The particle size was determined by Nanoparticle Tracking Analysis (NTA), and DLS. The UCLs were found to have a mean particle size of 100.6±1.1 with NTA and 100.7±0.9 with DLS, while ULC-211 had a mean particle size of 127±1.3 nm measured by DLS, while it was found to be around 108.1±0.6 nm using NTA **(Figure 1.a,b and Figure S2.a)**. The increase in the vesicle size upon miR211-5p complexation with UCLs could potentially be attributed to the fact that the complexation was performed after the downsizing and dialysis of the UCLs. This difference in results could be attributed to the dissimilar sizing principles used by the two techniques. While both DLS and NTA assess the hydrodynamic size of nanoparticles, the DLS simultaneously analyzes the ensemble of nanoparticles and becomes less accurate as the size variation within the sample (polydispersity) increases. On the other hand, NTA visualizes individual nanoparticles by the light they scatter, infers their sizes one by one from the observed motion [52, 53], and estimates the concentration of different size fractions. The measured concentrations of UCL and UCL-211 samples were 1 × 10^14^ and 1.68 × 10^14^ particles/mL, respectively **(Figure 1.b; Supplementary Figure S2.b includes a video illustrating the captured nanoparticle motion used as primary data for the NTA analysis)**. The standard deviations for concentration measurements were quite low. We also observed a reduction in the zeta potential upon UCL-211 complexation (from 51.25 mV before complexation to 42.3 mV), which was expected as the complexation between negatively charged miR211-5p neutralized the positive surface charges of UCLs. Since the zeta potential was still over +30mV, the suspension remained stable.

**Figure 1.**
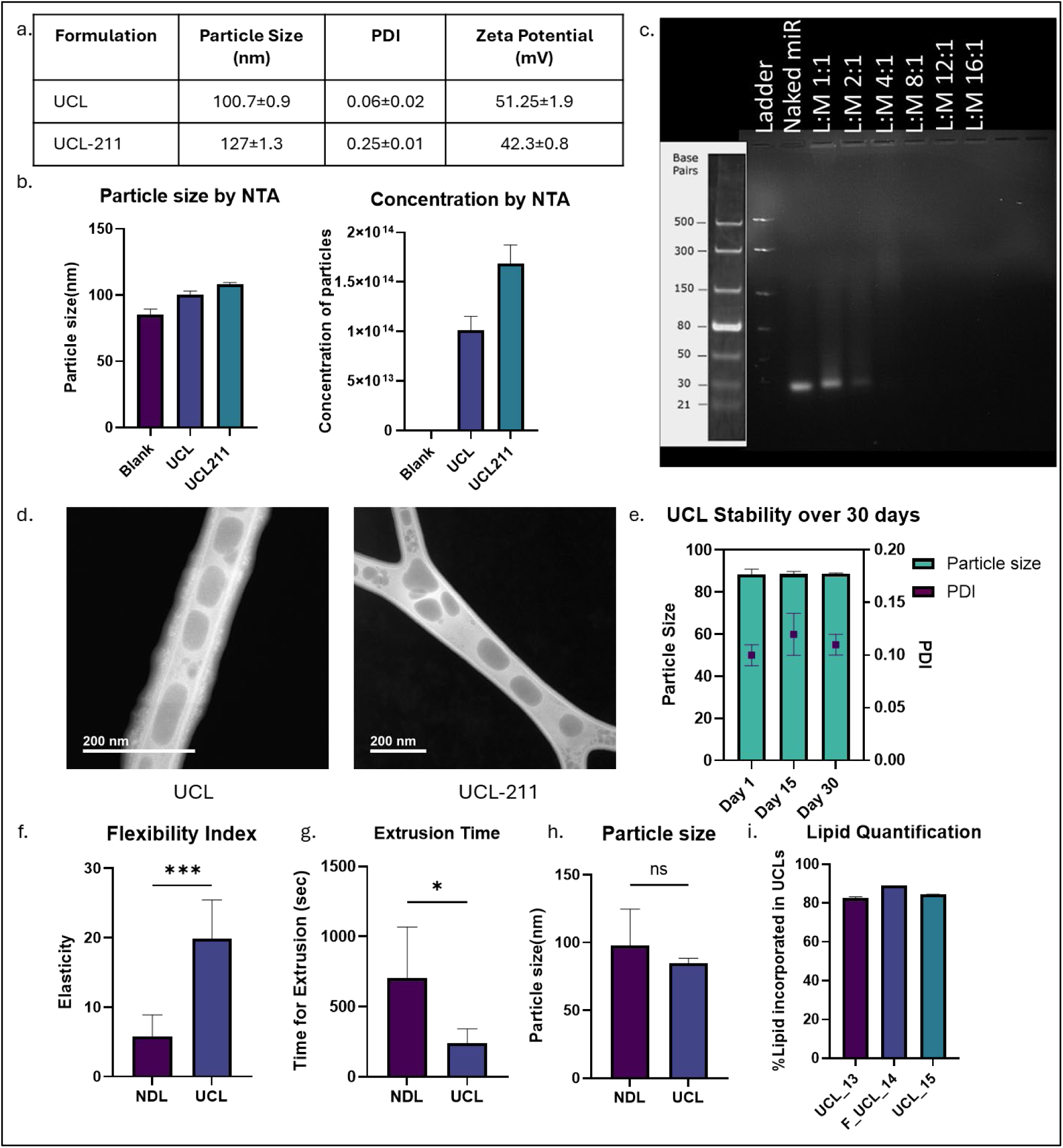
UCL and UCL-211 characterization. **a.** Particle size (nm), PDI and zeta potential measurement for UCL and UCL-211 using dynamic light scattering (DLS). **b.** Particle size and concentration determination by nanoparticle tracking analysis (NTA). **c.** Gel retardation assay of liposome (L) – miRNA (M) complexes. DOTAP: NaChol (8:1) was complexed with miR211-5-at different ratios ranging from 1:1 to 16:1 (w: w) and visualized for miRNA mobility on the agarose gel. Free miR211-5p was used as control. **d.** Transmission Electron Microscopy (TEM) images for UCL and UCL-211. **e.** Colloidal stability of UCL’s determined by particle size (nm) and PDI over 30 days. **f.** Flexibility index showing membrane elasticity for UCLs and NDLs. **g.** Time of extrusion of UCLs or NDLs at 100psi using a 50 nm membrane to evaluate deformability. **h.** Change in particle size after extrusion of UCLs or NDLs at 100psi using a 50 nm membrane to evaluate deformability (ns, not significant). **i.** Lipid quantification of three different batches of UCLs-UCL_13, F_UCL-14, and UCL_15 using Stewert assay. Data are reported as means ± S.D (n=3-6).

To further confirm efficient complexation between miR211-5p and UCLs, we performed gel retardation assay. UCLs (L) were complexed with 40µM miR211-5p (M) at different ratios ranging from 1:1 to 16:1 (v/v). The samples were then visualized for miRNA mobility across an agarose gel. In this assay, the UCL-211 complexes remained in the well, while free microRNAs migrated through the gel due to size and charge differences. As expected, while free miR211-5p migrated down the gel, for the UCL-211 complexes at UCL: miR211-5p (L:M) ratios higher than 4:1, no band corresponding to free unbound miR211-5p was observed. This confirmed that all miR211-5p molecules complexed with and neutralized by the positively charged UCLs **(Figure 1.c)**. Here we considered the L:M 1:1 as a negative control [27]. The morphology of the particles was determined by performing TEM imaging of UCL and UCL-211 formulations. It showed the UCLs had a particle size of around 100 nm which was similar to what was obtained using DLS and NTA **(Figure 1.d)**. Lacey carbon grids were used to perform TEM in this study, and we observed UCL deposition on the high-carbon portion of the grids. Some deformation of the UCLs to fit into the lacey part was observed, consistent with, but not conclusive of, some degree of elasticity.

We assessed the colloidal stability of the synthesized UCLs for over a period of 30 days (D30). We measured the particle size and PDI of these UCLs at D15 and D30. We did not observe any significant changes in particle size and PDI **(Figure 1.e)**.

We also sought to measure the amount of lipids that were incorporated into UCLs to confirm that most of the lipids initially added formed liposomes and were not lost. After the synthesis of UCLs, dialysis was performed for 24 h using 50kDa cut-off membranes to ensure the removal of any free lipids in the colloidal suspension. The percentage of lipids incorporated in the UCL preparations was then determined by Stewart Assay [44, 45] for three different UCL batches. Results showed an over 80% incorporation of the lipids into the UCLs **(Figure 1.i)**. This confirmed that over 80% of the lipids which were initially added had been incorporated into liposomes during UCL synthesis. The membrane elasticity for the UCLs was confirmed by performing deformability tests on UCLs as compared to conventional non-deformable liposomes (NDL). We synthesized NDLs using cholesterol instead of NaChol, thereby removing the edge activating agent, that conferred the deformability characteristics to liposomes, from the formulations. To assess the deformability, the UCL and NDLs were extruded through 50nm membranes at 100 psi (0.68 mPa) using a liposome extruder. We then calculated flux and membrane elasticity (flexibility index) by using previously reported equations [46]. We used the Stewart assay to determine the amount of DOTAP for each batch extruded. The calculated flexibility index showed that the UCLs had significantly more membrane elasticity than the NDLs **(Figure 1.f)**. The experiment was performed using two separately synthesized batches for both UCL and NDLs. The extrusion time for UCLs and NDLs was measured (in seconds) based on the time the formulations took to completely pass through the membrane, from the point where pressure was applied. UCLs took significantly less time to pass through the extruder compared to the NDLs **(Figure 1.g)**. No significant difference was observed when particle size was compared between UCLs and NDLs, which was measured using DLS **(Figure 1.h)**. However, PDI for NDLs was much higher compared to the UCLs.

We further measured the encapsulation efficiency of miR211-5p in UCLs using the Quant-iT™ microRNA Assay Kit. UCL-211 were mixed with 0.5% Triton-X100 and sonicated to break the liposomes to perform the assay. The encapsulation efficiency was 75.36±10.94%. Overall, these studies confirmed that the UCL formulations (UCL, fluorescently labeled UCL and UCL-211) were around 100nm in diameter, deformable, and stable with a high encapsulation efficiency.

### 3.3 Assessment of cytocompatibility and delivery of cargo *in vitro*

To ensure the cytocompatibility of our UCL-211 formulations, we assessed their potential effects on the viability of different primary human skin cells using the CCK-8 cytotoxicity assay. The viability of NHM and HDF cells in culture was tested after four hours of treatment with control (HEPES buffer 20mM), UCL-NC (negative miRNA control), and UCL-211 formulations. Cells treated with both UCL-NC and UCL-211 showed over 80% viability in comparison to the control. According to the International Organization for Standardization (ISO 10993-5:2009) guidelines, a topical formulation is considered cytocompatible if it maintains cell viability above 70% [54]. This high viability rate confirms the cytocompatibility of the tested formulations **(Figure 2.a, b)**.

**Figure 2.**
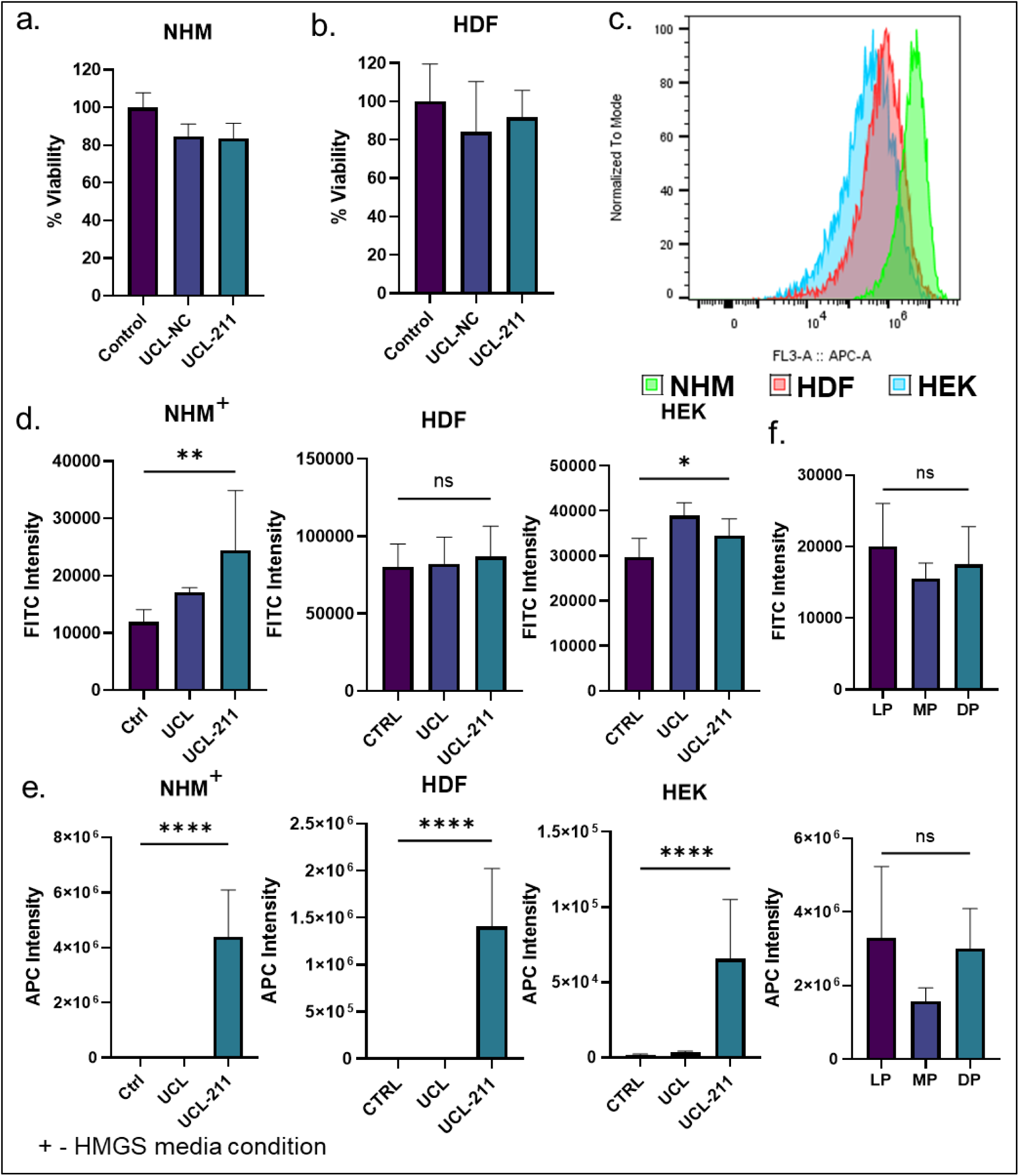
Cytotoxicity and uptake by primary cell cultures. CCK-8 assay confirmed UCL, UCL-NC, and UCL-211 were not cytotoxic to NHM **(a)** and HDF **(b)**. **c.** Histogram for APC intensity comparison of UCL-211 uptake in NHM, HDF and HEK (corresponding to miR211-5p conjugated with Cy5). **d.** Uptake of UCLs in NHM, HDF and HEK-FITC intensity (from F-DOTAP in UCLs). **e.** Uptake of UCLs in NHM, HDF and HEK-APC intensity (from Cy5 conjugated on miR211-5p). **f.** Pigmentation effect on uptake in light-pigmented (LP), medium pigmented (MP), and dark-pigmented (DP) melanocytes cultured in HMGS2 media. Data are represented as mean ± SD. (* p < 0.05; ** p < 0.01; *** p < 0.001; **** p < 0.0001; ns-not significant). Biological replicates (*N* = 2) and technical replicates (*n* = 3).

To evaluate the efficacy of UCL-211 in transporting its cargo (miR211-5p) into target melanoma cells, we compared the cellular uptake of free miR211-5p versus UCL-complexed miR211-5p (in UCL-211). To that end, we made fluorescently labeled UCL formulations (tagged with NBD (green) fluorescent dye) incorporating Cy5-labelled miR211-5p (pink), enabling us to quantify the cellular miRNA using flow cytometry. Initially, we treated NHM cells in *in vitro* cultures with fluorescently labeled free miR211-5p, empty UCLs and UCL-211, and performed flow cytometry using APC channel for the detection of Cy5 labeling on miRNAs, and FITC channel for the NBD labeling on the UCLs **(Supplementary Figure S3)**. The results from this study confirmed the uptake of UCL and UCL-211 based on the FITC intensity **(Supplementary Figure S3.a)**. As expected, a significant increase in APC intensity was observed when cells were treated with UCL-211 while no uptake was observed upon treatment with free miR211-5p, suggesting that UCL-211 efficiently delivered miR211-5p into NHM cells **(Supplementary Figure S3.b)**.

We then compared uptake in additional experiments with NHM (in HMGS2 media), HEK, and HDF cells in *in vitro* cultures and treated them with UCLs or UCL-211 complexes. NHM cells showed the highest uptake of complexed miR211-5p revealed by largest shift in the Cy5 intensity in a comparative histogram of uptake of UCL-211 between all three primary cell lines **(Figure 2.c)**. This observation is consistent with prior studies that report melanocytes have higher lipid uptake than other primary skin cells [48]. All cells showed an increase in the FITC signal upon treatment with fluorescently labeled UCL and UCL-211, indicating successful uptake of liposomes by cells **(Figure 2.d)**. However, the increase was not very prominent as F-DOTAP was incorporated in the UCLs at a very low concentration (1:100-F:DOTAP:DOTAP). Quantification of the APC intensity further confirmed the high accumulation of miR211-5p in other cells only when delivered by UCLs **(Figure 2.e)**.

We also evaluated the role of pigmentation and differentiation status on the uptake of UCLs. Melanocytes exist in several differentiation states in human skin and exhibit different levels of pigmentation (melanin) correlating to skin tone [47]. Both pigmentation level and differentiation status can affect the biological functions of melanocytes and the resistance to the therapeutic effects of small molecules [55–58]. Different culture media conditions can induce the differentiation or dedifferentiation of the NHM cells *in vitro*. Specifically, the presence or absence of protein kinase C activator-TPA (tetradecanoylphorbol acetate) influences NHM differentiation status in culture [6]. In the absence of TPA (i.e., HMGS2 media) NHMs are in a less pigmented melanocyte progenitor state while in the presence of TPA (i.e., HMGS media), NHM cells exhibit a differentiated and more pigmented phenotype [6].

To determine if the pigmentation level (melanin content) or differentiation status affected the uptake of UCL-211 by NHM cells, we performed comparative uptake studies across NHM cells with these variations. We used primary NHM cells isolated from the skin with either light pigmentation (LP), medium pigmentation (MP), or dark pigmentation (DP), in both HMGS2 (dedifferentiating) media conditions **(Supplementary Figures S3.c, d, e, f, g, h)** and HMGS (differentiating) **(Supplementary Figures S3.i, j, k, l)**. The LP NHMs are expected to exhibit the least amount of melanin while DP NHMs are expected to exhibit the highest. The uptake experiments revealed that the pigmentation level of NHM cells grown in HMGS2 media did not affect the UCL uptake, or UCL-211 accumulation in cells (measured by FITC and APC signal intensity, respectively) **(Figure 2.f, up and down)**. Thus, the melanin content of cells did not seem to interfere with the uptake of UCL-211 in dedifferentiating culture conditions. In contrast, in medium-pigmented (MP) NHM cells grown in HMGS media, a relatively small but consistent and significant decrease in uptake was observed **(Supplementary Figure S3.m, n)**. This decrease was not consistent across all differentiation conditions and did not correlate with pigment levels, making the reason for the relatively lower uptake observed only in MP melanocytes under differentiating conditions unclear. However, this could be attributed to the specific primary cell preparation procedures. Nonetheless, these studies confirmed the uptake of UCLs was more prominent in all NHM conditions and remained consistent across most differentiation conditions and pigmentation levels.

We further confirmed the results obtained in flow cytometry, by imaging the cells after uptake experiments using confocal microscopy. All three cell types, NHM, HDF and HEK, were treated with fluorescently labeled UCLs or UCL-211 following the same protocol as in flow cytometry experiments, and then visualized by fluorescent microscopy. We could observe the green and pink signals corresponding to UCLs (green) and miR-211-Cy5 (pink) in the cells treated with UCL-211, confirming the accumulation of labeled UCLs and miRNA in cells. The highest Cy5 intensity was observed in melanocytes compared to the other two primary cell cultures **(Figure 3.a, b, c; Supplementary Figure S4.a)**. Live cell imaging of NHM cells over 4.5 hours, and measurement of integrated intensity levels further confirmed the presence of higher Cy5 intensity (measured as the mCherry intensity) in UCL-211 treated groups compared to controls **(Supplementary Figure S4.b, c)**.

**Figure 3.**
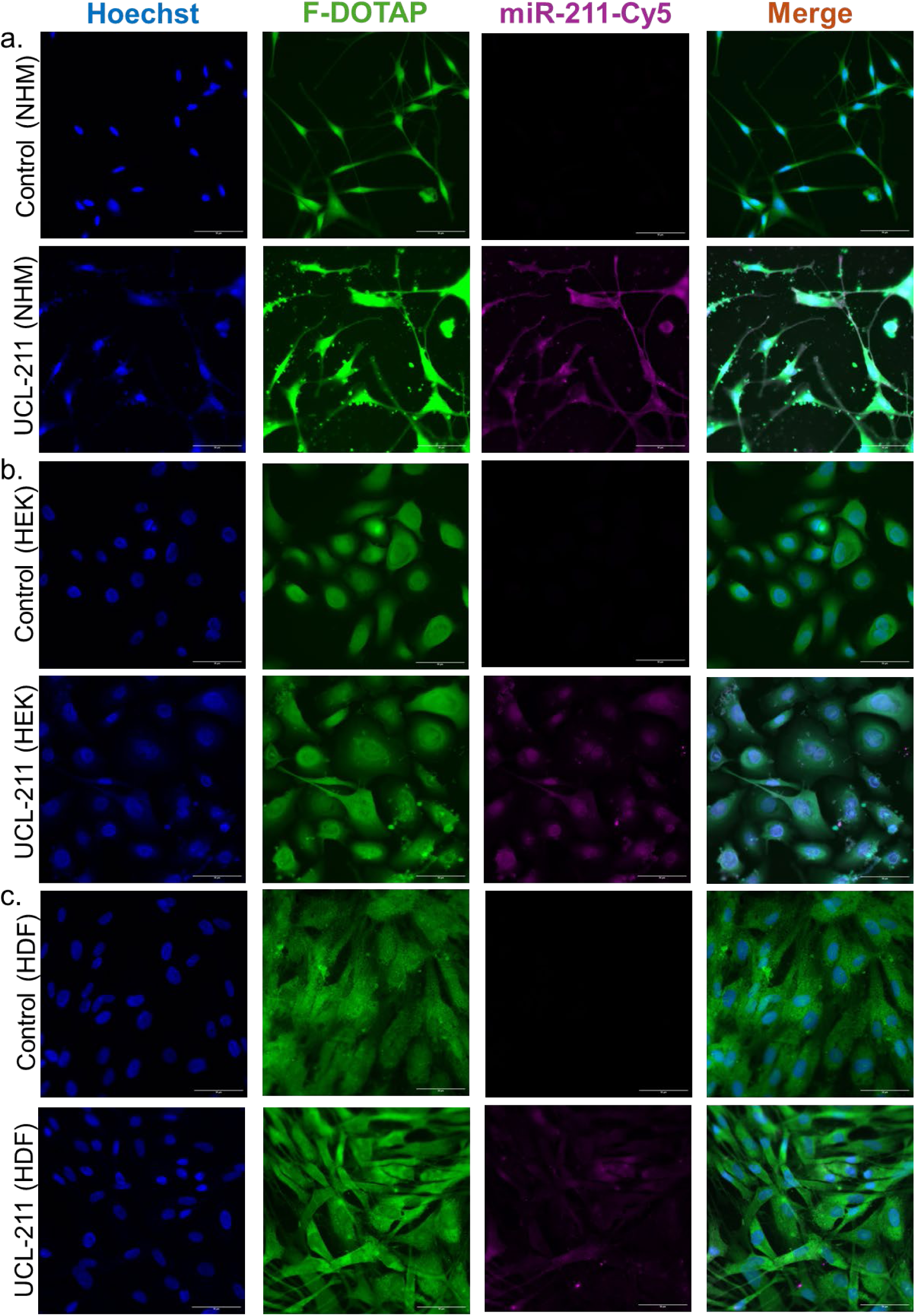
Fluorescent images for cellular uptake of UCL-211. **a.** NHM treated with HEPES 20mM (control) and UCL-211, **b.** HEK treated with HEPES 20mM (control) and UCL-211, **c.** HDFs treated with HEPES 20mM (control) and UCL-211. Blue-Hoechst, Green-UCL (F-DOTAP) and autofluorescence (inherently from cells), and Pink-miR211-5p-Cy5 (miR-Cy5).

To precisely evaluate the efficiency of miR211-5p delivery by UCL-211 liposomes, we used absolute quantitative PCR (qPCR), that is a highly sensitive and specific technique for nucleic acid quantification. We measured the intracellular accumulation of miR211-5p in NHM cells following treatment with our UCL-211 formulations, compared to UCL-NC negative controls. Since previous studies have demonstrated that growth in HMGS media upregulates the expression of endogenous miR211-5p to levels sufficient to drive the differentiation phenotype [6], we included non-treated HMGS-grown and HMGS2-grown NHM as additional controls for basal level of miR211-5p. To accurately assess the intracellular accumulation of miR211-5p delivered by UCL-211 liposomes and differentiate between internalized miRNA and any residual extracellular or surface-bound molecules, we selectively captured the cells using magnetic beads conjugated with CD117 antibody. We performed a rigorous washing step prior to elution, to eliminate any residual miR211-5p. The third wash solution (for UCL-NC and UCL-211) and final elution (obtained by detaching cells from magnetic beads) samples for all batches were then collected and miRNA copies per µL was quantified by absolute qPCR for miR211-5p (target miRNA) and miR-16-5p (endogenous control). The experiment was conducted in triplicate and elution 1,2,3 and wash 1,2,3 show results for each replicate.

As expected, wash solutions contained little miR211-5p and cells grown in HMGS media expressed more miR211-5p than their HMGS2-grown counterparts. The highest levels of intracellular miR211-5p were present in the UCL-211 treated group **(Figure 4.a)**, and the expression of the housekeeping miRNA (miR-16-5p) did not change across different conditions **(Figure 4.b)**. This confirmed that the results observed for UCL-211 were not false positive, but indeed associated with miR211-5p being delivered to the cells at levels that sufficiently match or exceed the endogenous levels known to trigger phenotype change.

**Figure 4.**
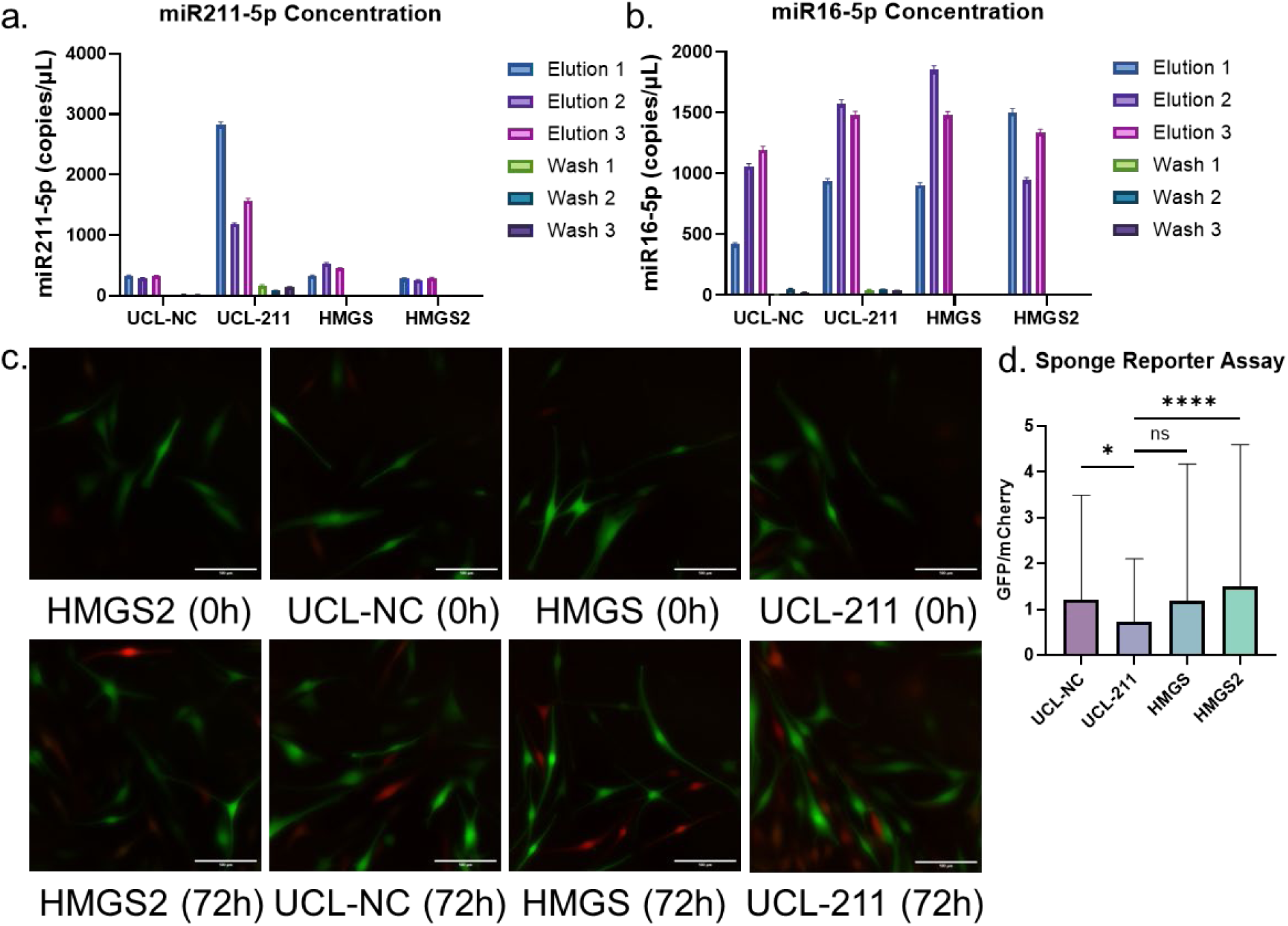
Analyzing biological functionality of miR211-5p delivered by the UCLs. Quantification of miR211-5p using Abs q-PCR. **a.** Concentration in copies/µL of miR211-5p in the total sample confirmed to be the highest levels in the UCL-211 treated group. **b.** Concentration in copies/µL of miR16-5p in the total sample. **c.** Live cell images from fluorescent protein-based miR211-5p sponge reporter assay with cells treated with HMGS2 (control), UCL-NC, HMGS and UCL-211 at 0h and 72h. **d.** Overall increase in the mCherry signal after 72 h and reduction in the GFP signal (calculated as GFP/mCherry ratio) (n=3). Data are represented as mean ± SD. (* p < 0.05; **** p < 0.0001; ns-not significant).

### 3.4 *In vitro* evaluation of the biological activity of UCL-delivered miR211-5p

The biological activity of delivered miR211-5p is a crucial aspect for its potential therapeutic application. We employed multiple orthogonal approaches to determine if internalized miR211-5p retains its functionality post-delivery. First, we used a previously described dual-fluorescent reporter system for miR211-5p activity assessment [6, 59]. The pLVX-anti-MIR211-5p reporter system detects active miR211-5p in cells, by the expression of two distinct fluorescent proteins as a measure for miR211-5p activity. The expression of ZsGreen fluorescent protein in pLVX-anti-miR211-5p construct is negatively regulated by the active miR211-5p, and the expression of mCherry fluorescent protein is positively correlated with the level of active miR211-5p [6]. Therefore, cells harboring pLVX-anti-MIR211-5p reporter appear mostly green under fluorescence microscopy in the absence of active miR211-5p, and red in its presence. We infected NHMs with lentiviral constructs expressing pLVX-anti-MIR211-5p reporter, and sorted cells expressing the biosensor within its linear range **(Supplementary Figure S5.a, b, c, d, e)**. The sorted cells then were put back into culture and treated with UCL formulations, similar to previous experiments. The cells were washed after 6h and the plates were placed on the PhaseFocus Livecyte QPI system for 72h, to quantify the ZsGreen (which were measured as the GFP levels) and mCherry levels. A shift towards red fluorescence in cells is an indicator of increased activity of miR211-5p in cells. Higher number of red (mCherry-positive) cells detected in cultures with HMGS media condition, consistent with prior literature that this condition induces endogenous miR211-5p expression **(Figure 4.c)**. Cells treated with UCL-211 exhibited a shift from green to red fluorescence, confirming the delivery of active miR211-5p by UCL-12 into cells **(Figure 4.c; Supplementary Figure S5.f, video)**. Control cells grown in HMGS2 condition or treated with UCL-NC did not exhibit a shift from the green to red fluorescent signal. The shift from the green to red fluorescence intensity was further quantified using ratiometric analysis on FIJI. We observed an overall increase in the mCherry signal and calculated the ratio of GFP to mCherry signal as a quantitative measure for miR211-5p activity. The GFP/mCherry ratio was significantly lower in the UCL-211 treated group than in the UCL-NC and Control (HMGS2) **(Figure 4.d)**.

Having demonstrated the delivery of biologically active miR211-5p mimic at levels equivalent or surpassing endogenous induction of the miRNA by HMGS, to NHM cells by UCL-211, we next asked whether UCL-211 exposure elicited the same phenotypic changes as exposure to HMGS did. Previous studies have shown that dedifferentiated NHMs (grown in HMGS2 media) will re-differentiate in HMGS media, both increasing miR211-5p expression levels and adopting a distinct morphology [6]. We, therefore, employed Phasefocus Livecyte® QPI live cell imaging to monitor changes in morphology.

Cells were grown in HMGS2 media and then treated with HEPES buffer (control), UCL-NC, and UCL-211. After 6h, the cells were washed, and then live imaged for 72h. At 0 h, the majority of cells were round and dendritic **(Figure 5.a, c, e, g)**. After 72 h, cells treated with UCL-NC or non-treated cells in HMGS2 media retained these morphologies **(Figure 5.d, h (right columns))**. In contrast, after 72 h, cells non-treated in HMGS media adopted the expected elongated morphology **(Figure 5.b (right column))**, a transition phenocopied by treatment with UCL-211 **(Figure 5.f (right column))**. Individual morphologic features were also tracked, such as median cell thickness **(Figure 5.i)**, median cell sphericity **(Figure 5.j)**, and median cell area **(Figure 5.k)**, and confirmed similarity between UCL-211 and HMGS conditions (leading to higher basal expression of miR211-5p in non-treated cells) for multiple parameters over the 72 h hours. Together, these experiments confirm that UCL-211 not only delivers intracellular miR211-5p but that the miRNA engages targets, inhibits translation, and induces cellular phenotypes associated with the endogenous microRNA.

**Figure 5.**
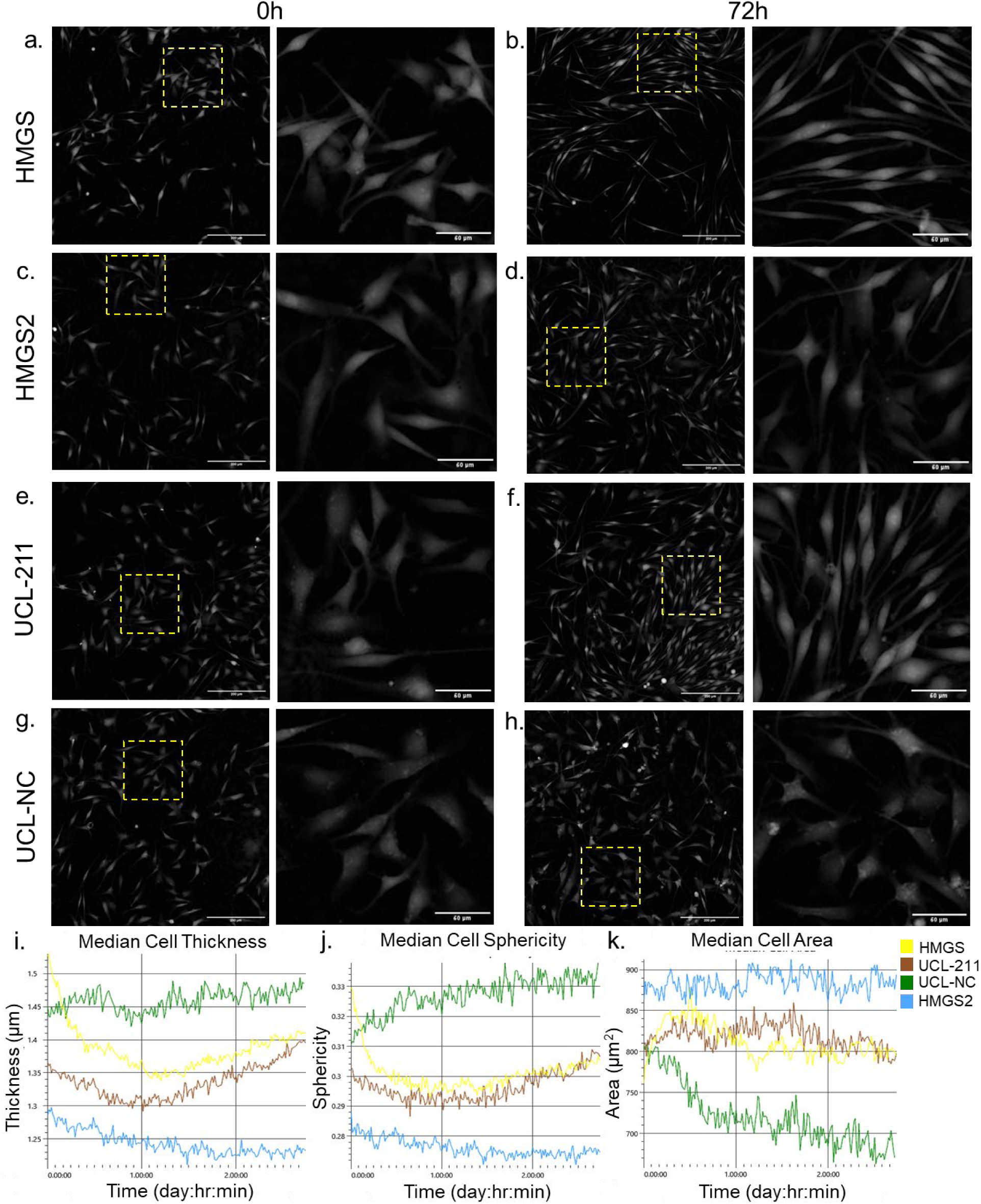
Increased miR211-5p levels causes morphological changes in NHM. Cells at time 0 h-HMGS (**a**), UCL-NC (**c**), UCL-211 (**e**) and HMGS2 (**g**). After 72 h treatment with HMGS media (positive control) **(b)** and UCL-211 **(f)**, cells became elongated due to the presence of miR211-5p. Cells treated with HMGS2 media (negative control) **(d),** and UCL-NC **(h)** were rounded in shape with no change. Integrated analysis for population morphology using Livecyte® Analyse software over 72 h confirmed the similarity between UCL-211 and HMGS conditions, including median cell thickness **(i)**, median cell sphericity **(j)**, and median cell area **(k)**.

### 3.5 *Ex vivo* study with skin tissue biopsies confirmed UCL-211 permeation

A permeability study using the Franz diffusion system was conducted to determine if the UCLs were able to permeate through intact skin and reach the epidermis-dermis junction, where the melanocytes reside. Skin from the same anatomical region of a healthy donor was used for this experiment, in order to exclude any variation in skin thickness. The samples were applied carefully on the skin exposed in the donor chamber (stratum corneum) and spread evenly. The tissue was collected after 24 hours, cleaned, fixed, and sectioned. Immunofluorescence staining was performed on the tissue sections for melan-A, a melanocyte-specific antibody. The presence of melanocytes was confirmed at the epidermis-dermis junction, which were stained red. In the tissue specimens treated with fluorescently labeled UCL-211, we observed liposomes around the epidermis which were not present in the control tissue samples, confirming that during the 24 h incubation time, the UCLs were able to permeate through the intact skin and cross the stratum corneum and travel through the epidermis **(Figure 6)**.

**Figure 6.**
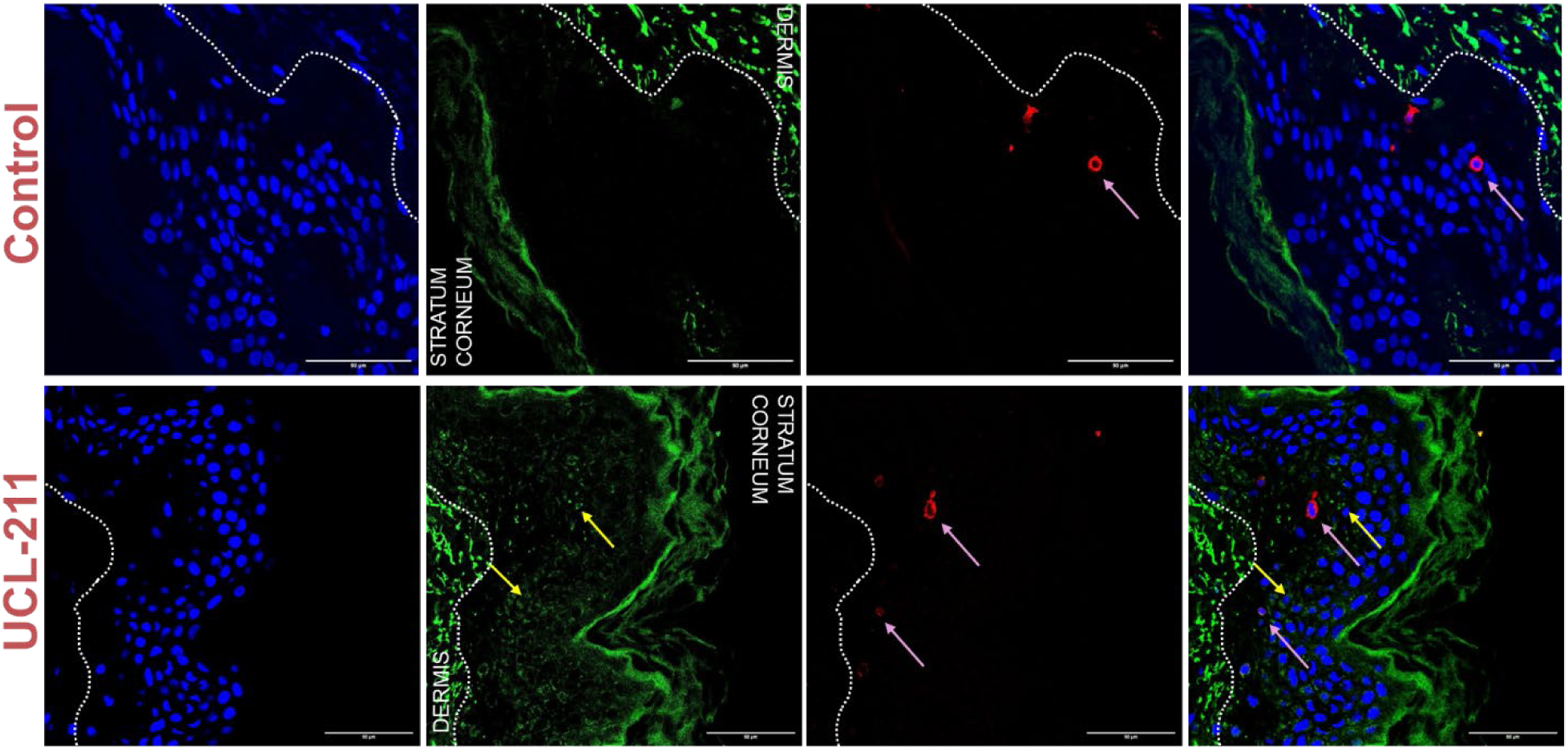
*Ex vivo* permeation of the UCLs. Treatments-Control (HEPES buffer 20mM) and UCL-211. Blue-Hoechst, Green-UCL (F-DOTAP) and autofluorescence, and Red-Melan-A (melanocyte marker). Yellow arrow: UCL-211. Pink Arrow: Melanocytes. A dotted line was drawn to separate the epidermis and dermis. Scale bar 50 μM.

### 3.6 *In vivo* evaluation shows UCL-211 treatment induces benign nevus formation from BRAFV600E melanocytes

After meticulous *in vitro* and *ex vivo* characterization of the UCL-211 delivery system, we evaluated the transdermal delivery of biologically active miR211-5p in a preclinical system using UCL-211. We employed an established and widely used genetically engineered mouse model (GEMM) of nevus formation [60]. The model selected harbors an inducible mutant BRAFV600E allele (BRAFCA) engineered into the endogenous BRAF locus, enabling temporal control of its expression. In the absence of Cre recombinase (Cre-ER), wild-type BRAF is expressed. However, upon transient expression of Cre-ER, BRAFV600E is activated and expressed. [49, 60]. These mice were bred to a line harboring a transgenic Cre-ER allele under the control of the *Tyrosinase* promoter – a melanocyte-enriched gene in skin [61]. The resulting transgenic animals enable the spatiotemporal control of BRAFV600E activation in tissues of interest. Direct topical application of 4-hydroxytamoxifen (4-HT) to the flanks of adult mice induces the formation of BRAFV600E melanocyte fields, some of which give rise to growth-arrested melanocytic-nevi over the course of 21 days with defined and reproducible kinetics [48]. We tested our formulations in this model by topically applying UCL-211, vemurafenib, UCL-NC (UCL complexed with non-targeting control miRNA), free miR211-5p mimic (diluted in HEPES buffer 20 mM), and NDL-211 (non-deformable liposome complexed with miR211-5p) to previously 4-HT-treated skin regions. The experimental design is summarized in **Figure 7.a**.

**Figure 7.**
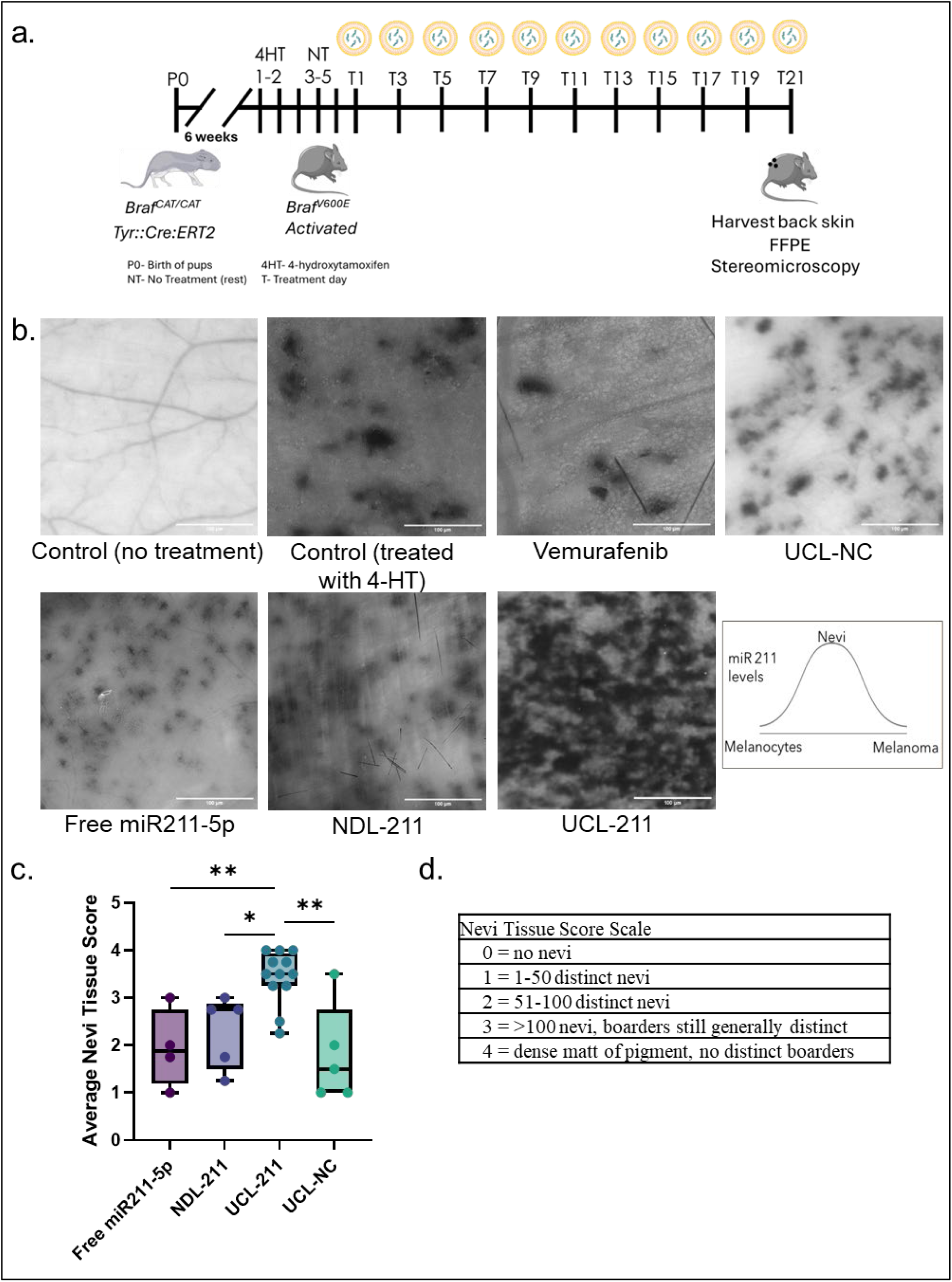
*In vivo* evaluation of the UCL-211 delivery system. **a.** Schematic for developing the transgenic nevus growth-arrested mouse model. **b.** Images of mouse skin on T (treatment day) 21-Control (no 4-HT application-no nevi), control (4-HT treatment-nevi), Vemurafenib (positive control), UCL-NC, free miR211-5p, NDL-211 and UCL-211. Also, a pictorial depiction showing an increase in miR211-5p expression in benign nevi upon miR211-5p treatment compared to healthy melanocytes and melanoma. **c.** Average nevi tissue score comparison between free-miR211-5p, NDL-211, UCL-211 and UCL-NC treated groups. **d.** Nevi tissue score scale, used for semi-quantitative analysis of the nevi. Data are represented as Box and whisker plot (min to max, showing all data points) (* *p <* 0.01; ** *p <* 0.001). Number of mice-UCL-NC (n=5), Free211-5p (n=4), NDL-211 (n=5) and UCL-211 (n=12).

Specifically, on the dorsal side of the skin that was previously treated with 4-HT, various treatments were applied every other day for 21 days. As expected, the application of 4-HT induced nevi, whereas no nevi were observed in the absence of BRAFV600E activation by 4-HT. Vemurafenib was a positive control as it is a selective BRAFV600E inhibitor and upon treatment it reduced nevus density, confirming that the nevi formation relied on the oncogene. Treatment with UCL-211 **(Supplementary Figure S6)**, but neither UCL-NC, NDL-211 **(Supplementary Figure S7),** nor free miR211-5p mimic **(Supplementary Figure S8)** resulted in a striking increase in the density of nevi on the 4-HT treated patches. Results obtained for UCL-211 were compared based on the delivery system used (free-miR211-5p and NDL-211) or to the non-targeting miRNA (UCL-NC) **(Figure 7.c)**, which was quantified using the nevi tissue score scale **(Figure 7.d)**. Previous study from our lab has demonstrated *in vitro,* that the level of miR211-5p was elevated in nevi melanocytes, compared to healthy melanocytes and melanoma [6] (**Figure 7.b** (pictorial depiction)). These findings demonstrate that biologically active miR211-5p was transported and delivered to melanocytes by the UCL-211 system. Our results also revealed that miR211-5p overexpression increased nevi formation from BRAFV600E+ melanocytes in UCL-211 treated zones, which were significantly higher compared to regions treated with UCL-NC **(Figure 7.b, c)**. Furthermore, this effective delivery of miR211-5p to melanocytes is dependent on the ultra-deformable nature of UCLs, as non-deformable liposomes (NDL-211) and free miR211-5p mimic failed to elicit the same biological phenotype.

## 4. Conclusions and future studies

Here, we reported the development and evaluation of a transdermal delivery system using ultradeformable cationic liposomes (UCLs) for topical delivery of miRNAs to melanocytes, present at the skin epidermal-dermal junction. The UCLs were characterized for their physicochemical properties and deformability using various approaches, and extensively evaluated for the uptake and cargo delivery. Our study demonstrated that the melanocytes have the highest rate of UCL uptake compared to keratinocytes and fibroblasts. The differentiation status and pigmentation level of melanocytes had negligible effect on the uptake of UCLs. Multiple orthogonal approaches revealed *in vitro* biological activity of miR211-5p mimics delivered by UCL-211 into cells. *Ex vivo* permeation study using human skin biopsies confirmed the permeation of UCL-211 across the epidermis. We further demonstrated the successful targeted delivery of miRNAs to melanocytes in an *in vivo* mouse model. These results expand upon previous observations of miR211-5p being enriched in melanocytic nevi, as compared to healthy or malignant melanocytes, and that the endogenous expression of miR211-5p is sufficient to drive the nevus-associated phenotypes in BRAFV600E human melanocytes *in vitro* [6]. This is the first report of the *in vivo* confirmation of the combination of BRAFV600E mutation and miR211-5p expression driving nevus formation. While expected, the observation is noteworthy, not only as the first demonstration of transdermal delivery of a miRNA to melanocytes *in vivo* eliciting a complex biological effect, but also as a confirmation that the mechanisms that underlie nevus formation can be manipulated to alter the dynamics of tumor formation, offering new avenues for dermatological interventions.

Our study successfully establishes that UCLs can be utilized for topical delivery of other miRNAs or siRNAs to melanocytes to alter their behavior. In future studies, this approach can be used to rapidly explore the utility of transdermal small RNA delivery to treat pigment-related skin conditions, disorders, and diseases.

## Funding

This work was supported by the Department of Defense Melanoma Research Program (W81XWH2010530 awarded to MWV and RLJ) and the National Cancer Institute (R01CA229896 to RLJ). We acknowledge the direct financial support for the research reported in this publication provided by seed grants from the Huntsman Cancer Institute Melanoma Research Center. This research was also supported by the University of Utah Graduate Research Fellowship awarded to TC (2023-24). We utilized the Shared Resources for Research Informatics and Biorepository and Molecular Pathology, each supported by the National Cancer Institute of the National Institutes of Health under Award Number P30CA042014.

## CRediT authorship contribution statement

**Tanya Chhibber:** Data curation, Formal analysis, Investigation, Methodology, Project administration, Resources, Visualization, Writing – original draft, **Michael T. Scherzer:** Data curation, Formal analysis, Investigation, Writing – review & editing, **Anastasia Prokofyeva:** Data curation, Formal analysis, Investigation, Writing – review & editing, **Carly Becker:** Data curation, Formal analysis, Investigation, Writing – review & editing, **Eric Smith:** Data curation, Methodology, Investigation, Writing – review & editing, **Rebecca Goldstein Zitnay:** Data curation, Formal analysis, Writing – review & editing, **Nitish Khurana:** Methodology, Writing – review & editing, **Dekker C. Deacon:** Supervision, Methodology, Writing – review & editing, **Mathew VanBrocklin:** Methodology, Resources, Funding acquisition, **Mikhail Skliar:** Data curation, Formal analysis, Investigation, Writing – review & editing, **Hamidreza Ghandehari*:** Conceptualization, Funding acquisition, Supervision, Writing – review & editing, Resources, **Robert Judson-Torres*:** Conceptualization, Funding acquisition, Supervision, Writing – review & editing, Resources, **Paris Jafari*:** Conceptualization, Funding acquisition, Supervision, Writing – review & editing, Resources.

## Competing interests

The authors have no competing interests to declare.

## Acknowledgments

We are grateful to the patients who consented to the use of discarded surgical tissue in these studies, and the staff of the Huntsman Cancer Institute Biorepository and Molecular Pathology Shared Resource who performed consenting and tissue collection. This work made use of Nanofab EMSAL shared facilities of the Micron Technology Foundation Inc. Microscopy Suite sponsored by the John and Marcia Price College of Engineering, Health Sciences Center, and The Office of the Vice President for Research. This work also used the University of Utah Nanofab shared facilities supported, in part, by the MRSEC Program of the NSF under Award No. DMR-112125″ We acknowledge the University of Utah Flow Cytometry Core facility, supported by the Office of the Director of the National Institutes of Health under Award Number S10OD026959 and the University of Utah Cell Imaging Core facility.

## Notes

### Competing Interest Statement

The authors have declared no competing interest.

