## Supplementary figures for "Transdermal Delivery of Ultradeformable Cationic Liposomes Complexed with miR211-5p (UCL-211) Stabilizes BRAFV600E+ Melanocytic Nevi"

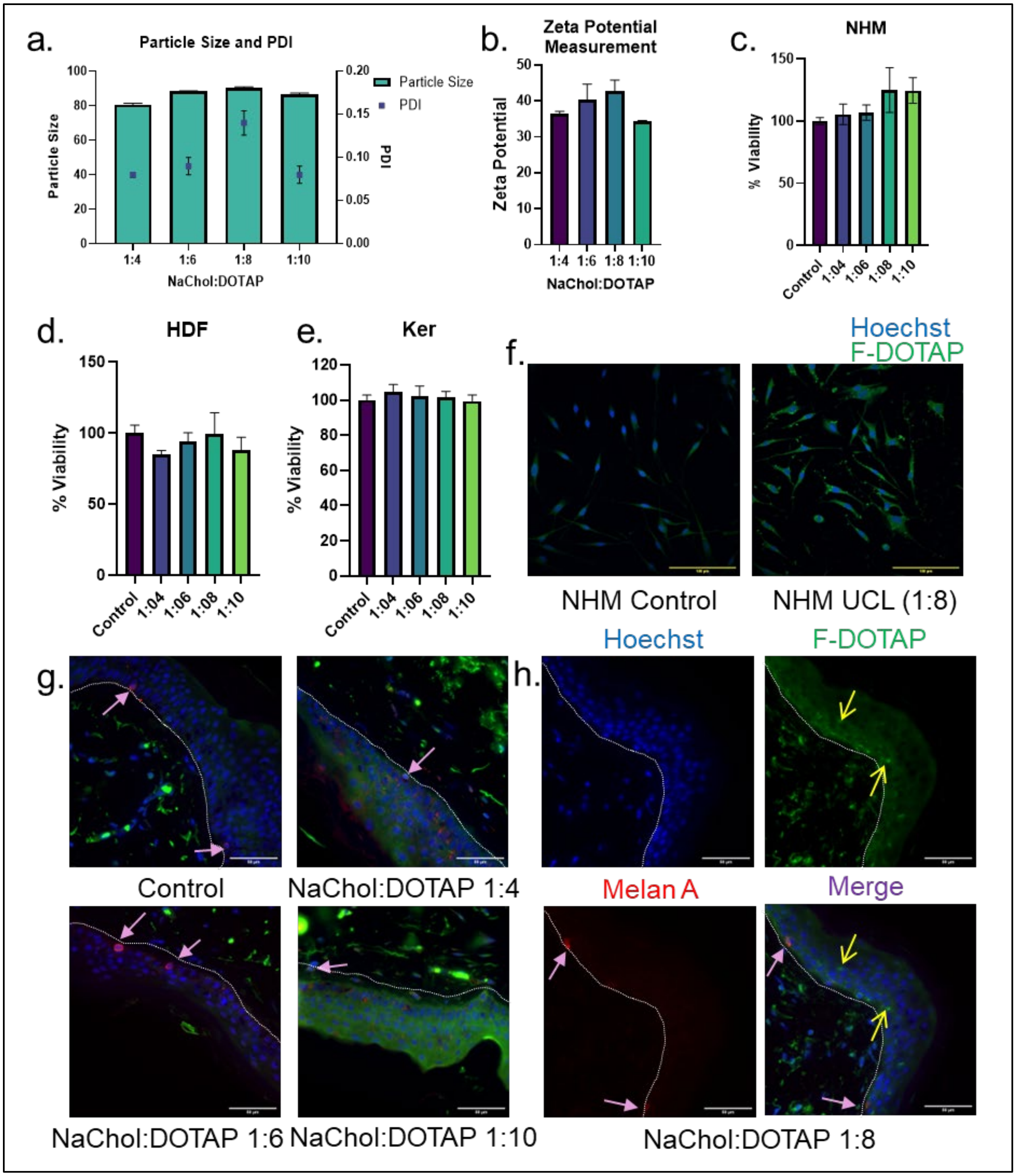


**Supplementary Figure 1. UCL characterization. (a)** Particle size (nm) and PDI measurement for NaChol: DOTAP 1:4, 1:6, 1:8 and 1:10 ratio using dynamic light scattering (DLS). (**b)** Zeta potential measurement for NaChol: DOTAP 1:4, 1:6, 1:8 and 1:10. Cytotoxicity studies with **(c)** normal human melanocytes (NHM), **(d)** human dermal fibroblasts (HDF) and **(e)** human keratinocytes (Ker) treated with UCLs with different NaChol: DOTAP ratios (1:4, 1:6, 1:8 and 1:10). (**f)** Uptake in NHM. Confocal microscopy images of NHM after 4 h treatment with HEPES buffer 20mM (control) and fluorescently labeled UCL 1:8. (**g)** Permeability studies with deidentified human skin tissue using Franz diffusion system. Confocal images (merged) for control, NaChol: DOTAP 1:4, 1:6, and 1:10 ratio. **(h)** Confocal images for NaChol: DOTAP 1:8. The nuclei were stained with Hoechst, F-DOTAP from the UCLs, melanocytes stained with melanocyte marker Melan A and merge images. Yellow arrows highlight the presence of liposomes in the epidermis layer in the NaChol: DOTAP 1:8 batch. Pink arrows highlight the presence of melanocytes in the tissue. A dotted line was drawn to separate the epidermis and dermis. Data are reported as means ± S.D. (n = 3).

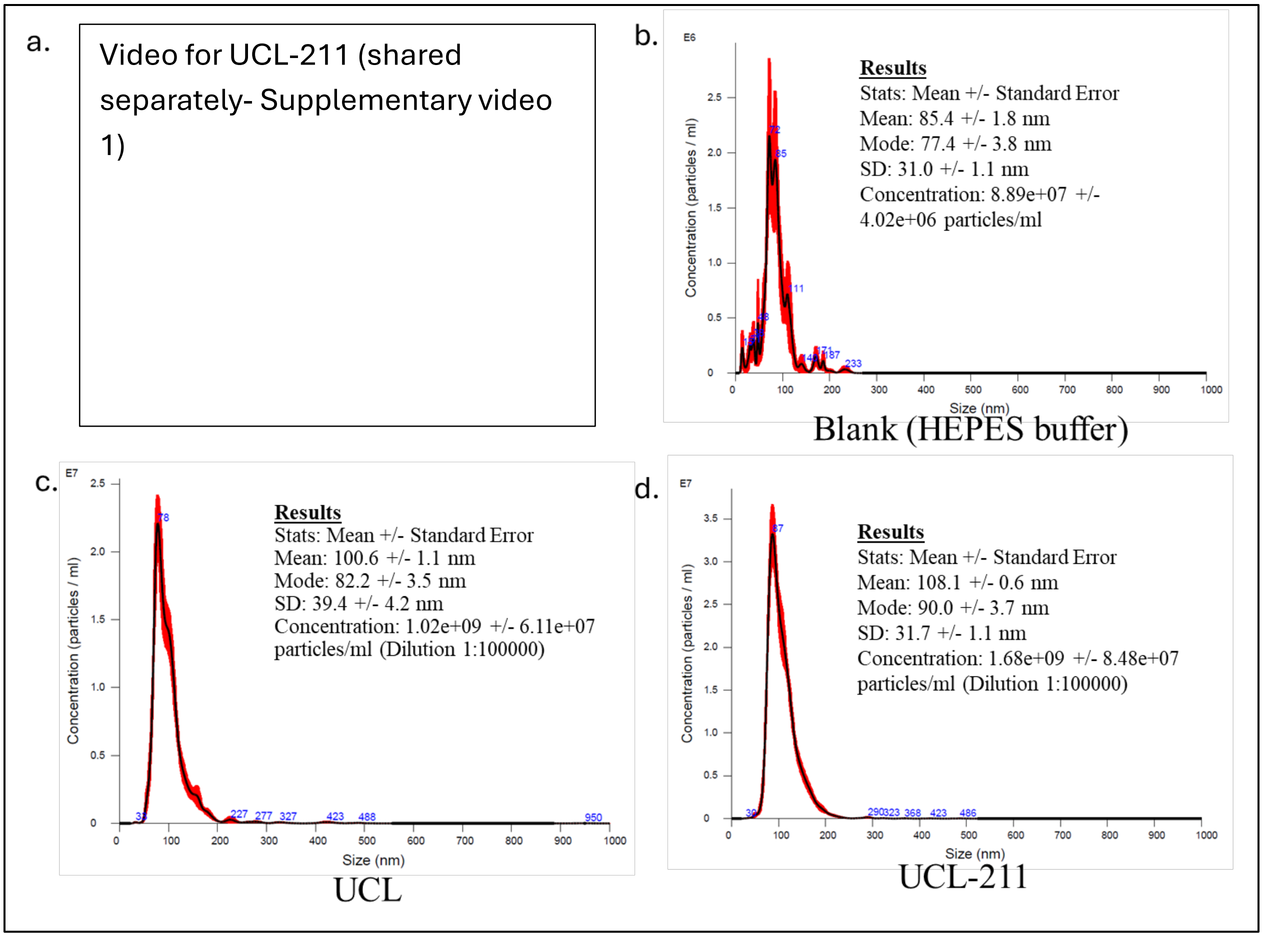


**Supplementary Figure 2: (a)** Video illustrating the captured nanoparticle motion used as primary data for the NTA analysis for UCL-211. NTA analysis for particle concentrations and size distributions in **(b)** HEPES buffer (Blank), **(c)** UCL and **(d)** UCL-211. Inserts list the statistics of the measured size distributions (mode and mean sizes and the standard deviation characterizing the width of the distribution) and overall particle concentrations for each sample. The red shading around the size distributions indicates ± SD of the concentration measurements for particles of different sizes.

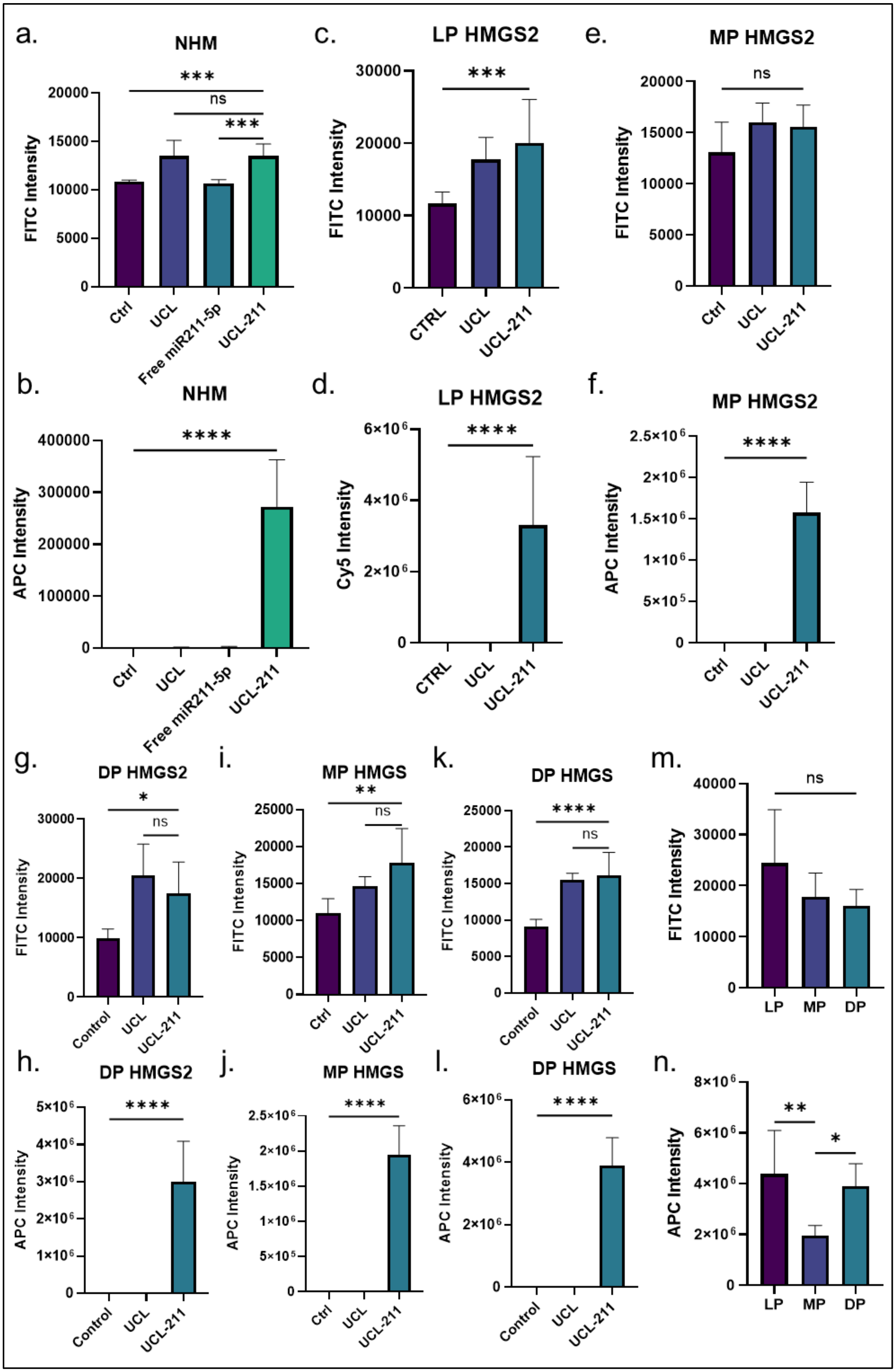

a.
NHM
FITC Intensity
Ctrl
UCL
Free miR211-5p
UCL-211
***
ns
***
b.
NHM
APC Intensity
****
c.
LP HMGS2
FITC Intensity
CTRL
UCL
UCL-211
***
d.
LP HMGS2
Cy5 Intensity
****
e.
MP HMGS2
FITC Intensity
ns
f.
MP HMGS2
APC Intensity
****
g.
DP HMGS2
FITC Intensity
Control
*
ns
h.
DP HMGS2
APC Intensity
****
i.
MP HMGS
FITC Intensity
**
ns
j.
MP HMGS
APC Intensity
****
k.
DP HMGS
FITC Intensity
****
ns
l.
DP HMGS
APC Intensity
****
m.
FITC Intensity
LP
MP
DP
ns
n.
APC Intensity
**
*

**Supplementary Figure 3**: UCL in melanocytes with different pigmentation levels. Comparison of uptake of UCL-211 and free miR211-5p in NHM**- (a)** FITC intensity (F_DOTAP on UCLs) and **(b)** APC (miR211-Cy5) intensity. Uptake of UCLs in light-pigmented melanocytes (LP) in HMGS2 media, **(c)** FITC intensity **(d)** APC intensity. Uptake of UCLs in medium pigmented melanocytes (MP) in HMGS2 **(e)** FITC intensity **(f)** APC intensity and HMGS **(i)** FITC intensity **(j)** APC intensity media conditions. Uptake of UCLs in dark pigmented melanocytes (DP02) in HMGS2 **(g)** FITC intensity **(h)** APC intensity and HMGS **(k)** FITC intensity **(l)** APC intensity media conditions. Comparison of effect of pigmentation in NHM for UCL and UCL-211 uptake in cells grown in HMGS media condition **(m)** FITC intensity **(n)** APC intensity.

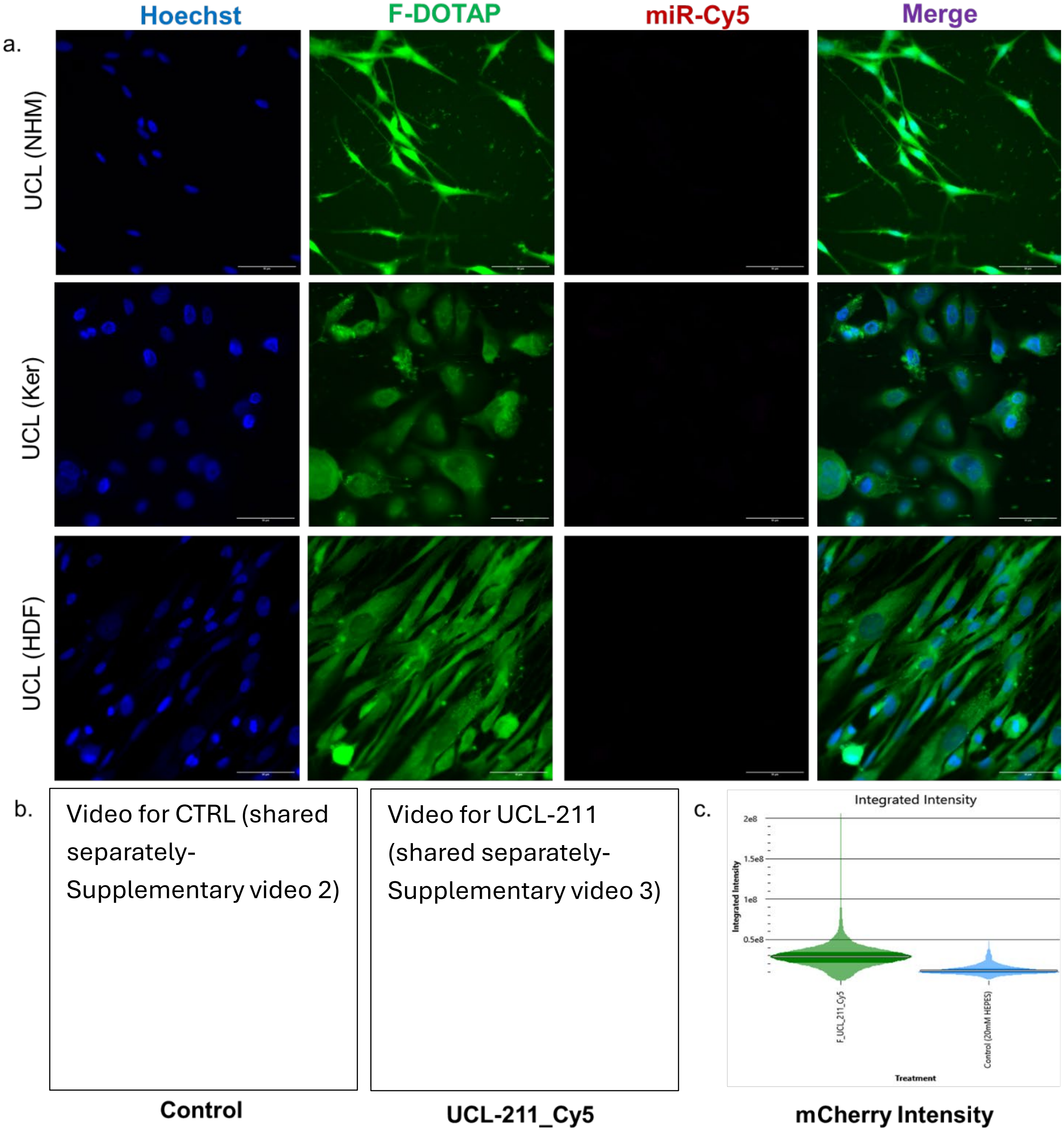


**Supplementary Figure 4**. a. Fluorescent images for cellular uptake of UCLs- Melanocytes (NHM), Keratinocytes (Ker) and HDFs. Blue- Hoechst, Green- UCL (F-DOTAP) and autofluorescence (inherently from cells), and Pink- miR211-5p-Cy5 (miR-Cy5). b. Videos of live imaging of NHM- Control and UCL-211_Cy5. c. Integrated mCherry intensity from Livecyte Analyse software.

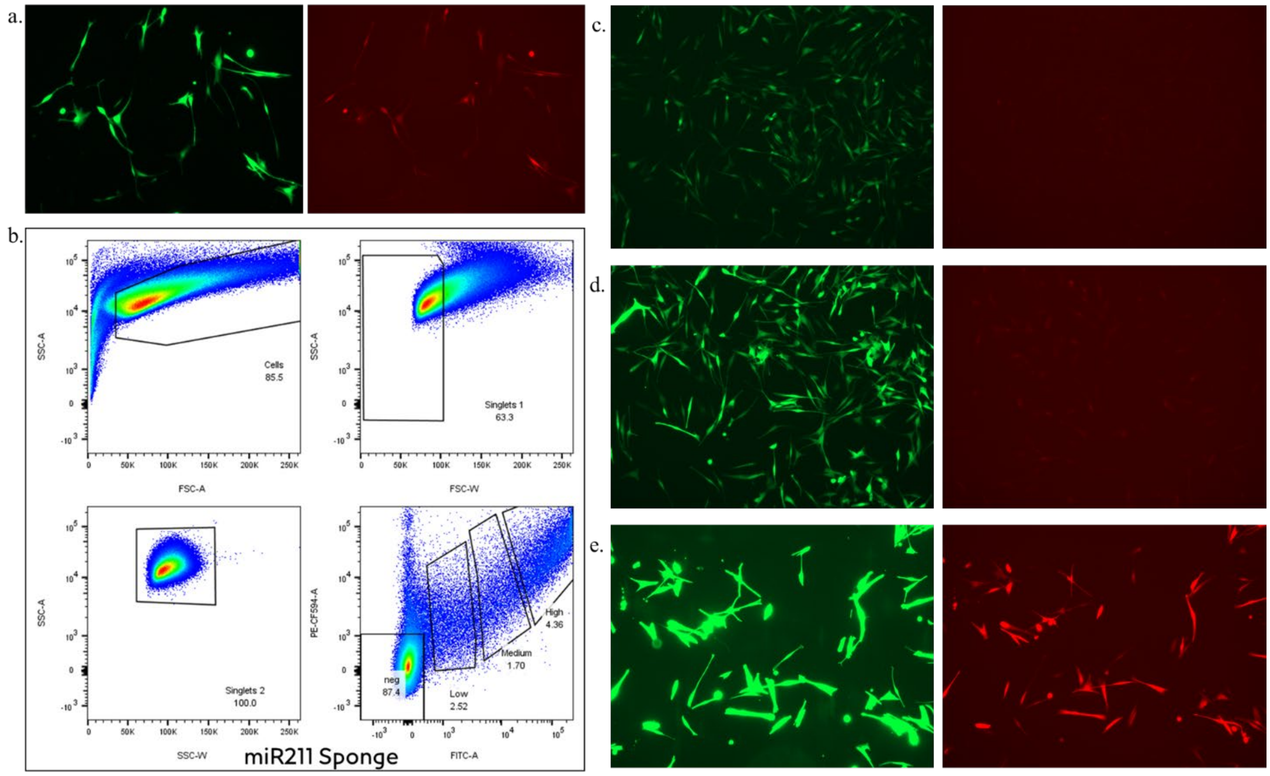


Videos for HMGS2 (shared separately- Supplementary video 4)

Videos for UCL-NC (shared separately- Supplementary video 5)

Video for HMGS (shared separately- Supplementary video 6)

Video for UCL-211 (shared separately- Supplementary video 7)

**Supplementary Figure 5**: Generation of fluorescent protein-based miR211-5p sponge reporter. **a.** NHM after transfection with the viral supernatant. **b.** Sorting of NHM with miR211 sponge into low, medium and high populations. **c.** NHM with low GFP (green)/ mCherry (red) reporter signal, **d.** NHM with medium GFP/mCherry reporter signal and **e. c.** NHM with high GFP/ mCherry reporter signal. **f.** Time-lapse videos from fluorescent protein-based miR211-5p sponge reporter assay with cells treated with HMGS2, UCL-NC, HMGS, and UCL-211. Green- GFP+, Red- mCherry+

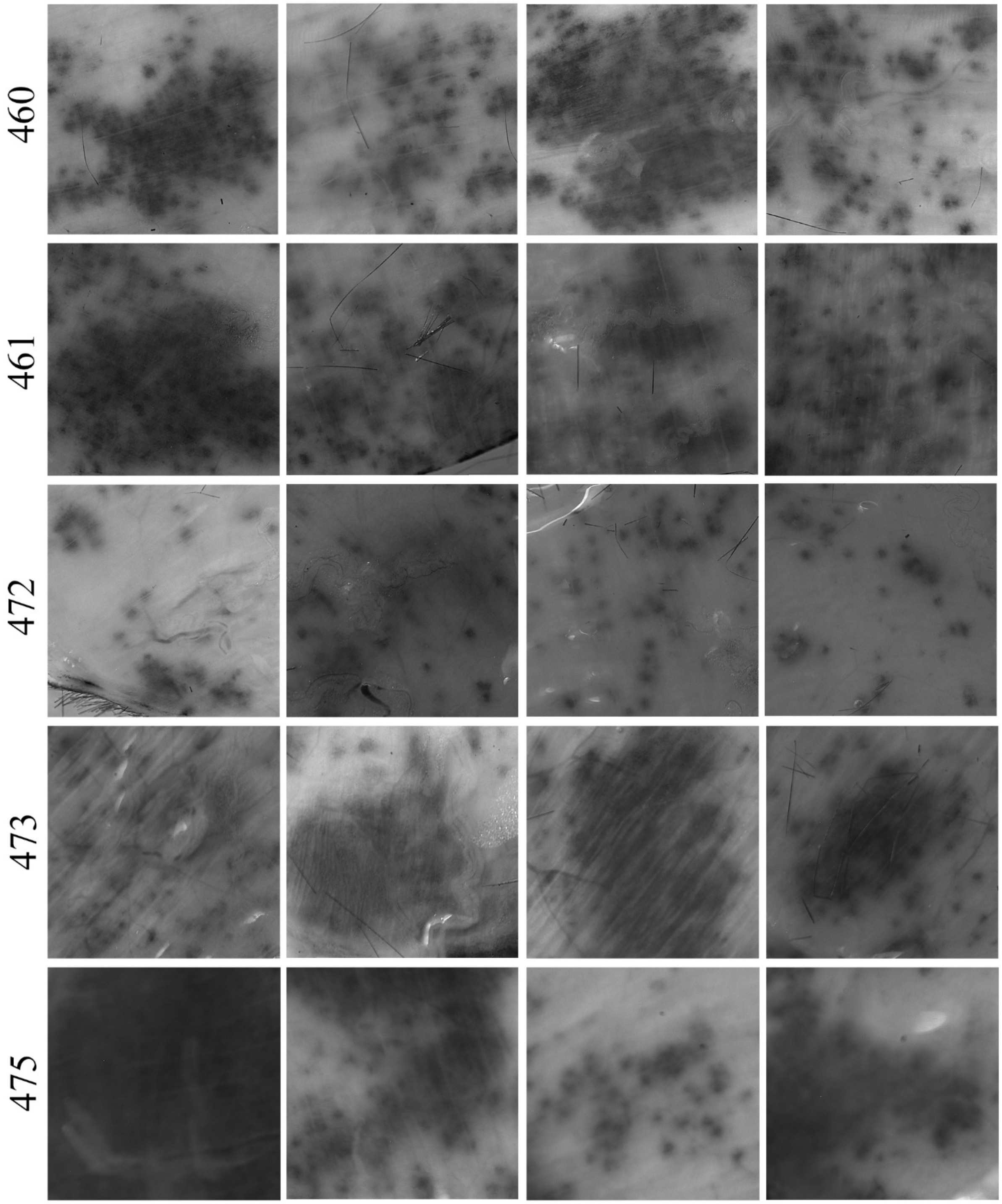


**Supplementary Figure 6**: Images of mice skin in the UCL-211 treated group considered per group for counting the number of nevi

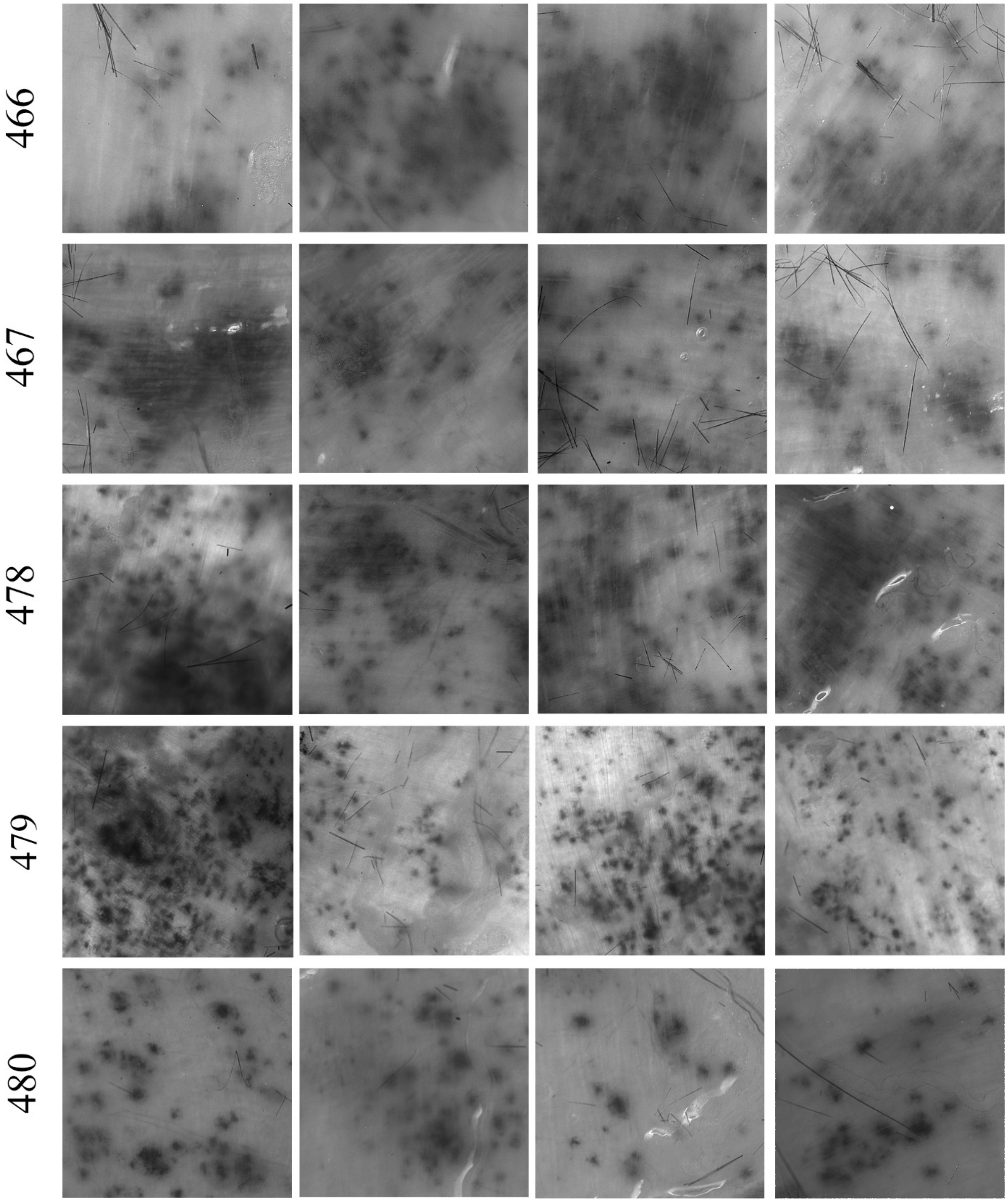


**Supplementary Figure 7**: Images of mice skin in the NDL-211 treated group considered per group for counting the number of nevi

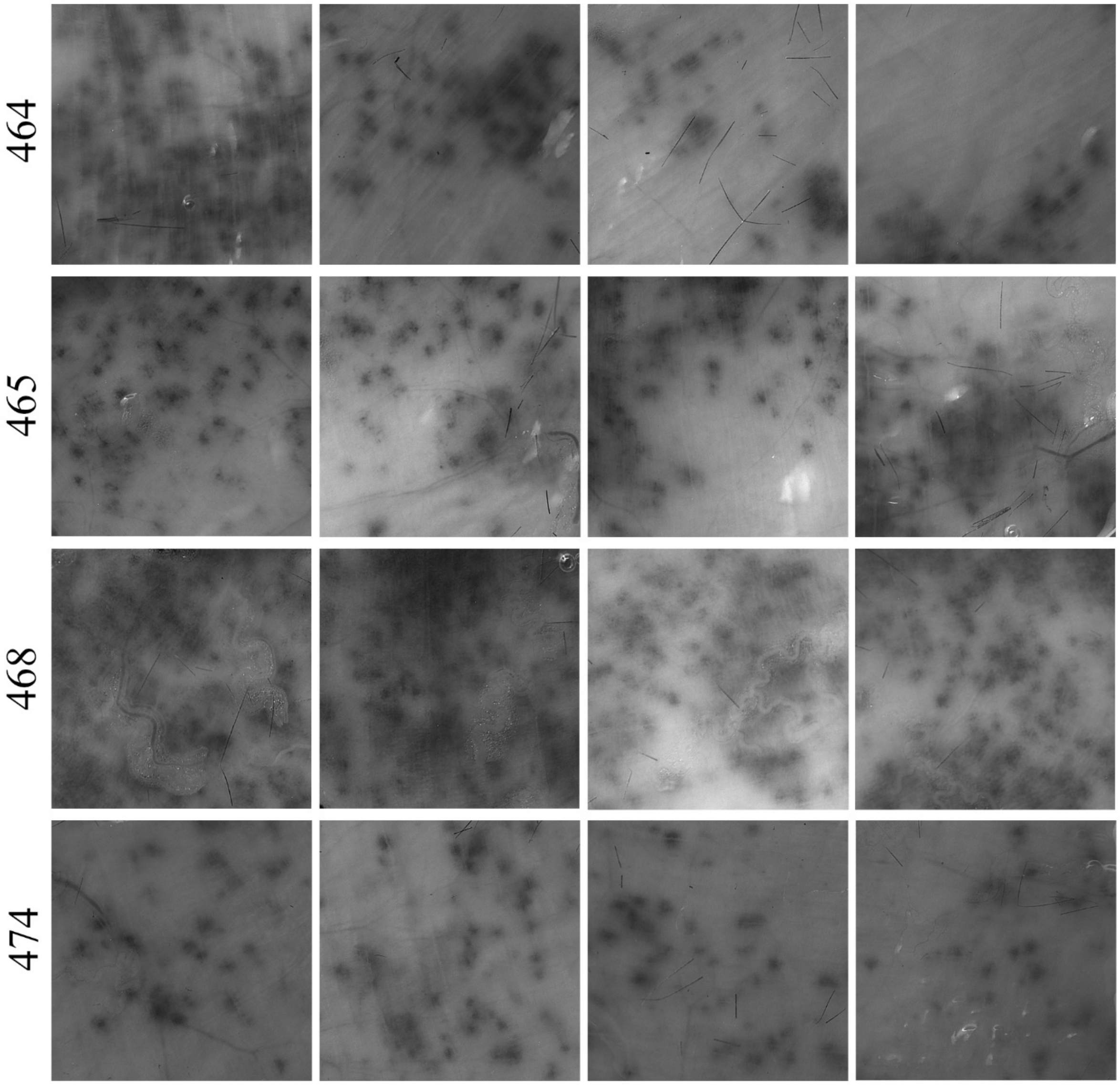


**Supplementary Figure 8**: Images of mice skin in free miR211-5p treated group considered per group for counting the number of nevi
